# Nanobodies against SARS-CoV-2 reduced virus load in the brain of challenged mice and neutralized Wuhan, Delta and Omicron Variants

**DOI:** 10.1101/2023.03.14.532528

**Authors:** María Florencia Pavan, Marina Bok, Rafael Betanzos San Juan, Juan Pablo Malito, Gisela Ariana Marcoppido, Diego Rafael Franco, Daniela Ayelen Militello, Juan Manuel Schammas, Sara Bari, Argentinian AntiCovid Consortium, William B. Stone, Krisangel López, Danielle L. Porier, John Muller, Albert J. Auguste, Lijuan Yuan, Andrés Wigdorovitz, Viviana Parreño, Lorena Itatí Ibañez

**Affiliations:** CONICET Universidad de Buenos Aires, Instituto de Química Física de los Materiales, Medio Ambiente y Energía (INQUIMAE).; Incuinta, Instituto Nacional de Tecnología Agropecuaria (INTA).; Instituto de Virología e Innovaciones Tecnológicas, Consejo Nacional de Investigaciones Científicas y Técnicas (IVIT-CONICET); Departamento de Química Biológica, Instituto de Química Biológica de la Facultad de Ciencias Exactas y Naturales (IQUIBICEN) CONICET, Ciudad Universitaria, Ciudad Autónoma de Buenos Aires, Buenos Aires, Argentina; Instituto de Investigación Patobiología, Centro de Investigaciones en Ciencias Veterinarias y Agronómicas (CICVyA), Instituto Nacional de Tecnología Agropecuaria (INTA).; Centro de Investigaciones en Ciencias Veterinarias y Agronómicas (CICVyA), Instituto Nacional de Tecnología Agropecuaria (INTA).; Department of Entomology, College of Agriculture and Life Sciences, Fralin Life Science Institute, Virginia Polytechnic Institute and State University, Blacksburg, USA; Center for Emerging, Zoonotic, and Arthropod-borne Pathogens, Virginia Polytechnic Institute and State University, Blacksburg, USA; Department of Biomedical Sciences and Pathobiology, Virginia-Maryland College of Veterinary Medicine, Virginia Polytechnic Institute and State University, Blacksburg, USA

## Abstract

In this work, we developed llama-derived nanobodies (Nbs) directed to the receptor binding domain (RBD) and other domains of the Spike (S) protein of SARS-CoV-2. Nanobodies were selected after the biopanning of two VHH-libraries, one of which was generated after the immunization of a llama (*lama glama*) with the bovine coronavirus (BCoV) Mebus, and another with the full-length pre-fused locked S protein (S-2P) and the RBD from the SARS-CoV-2 Wuhan strain (WT). Most of the neutralizing Nbs selected with either RBD or S-2P from SARS-CoV-2 were directed to RBD and were able to block S- 2P/ACE2 interaction. Three Nbs recognized the N-terminal domain (NTD) of the S-2P protein as measured by competition with biliverdin, while some non-neutralizing Nbs recognize epitopes in the S2 domain. One Nb from the BCoV immune library was directed to RBD but was non-neutralizing. Intranasal administration of Nbs induced protection ranging from 40% to 80% against COVID-19 death in k18-hACE2 mice challenged with the WT strain. Interestingly, protection was not only associated with a significant reduction of virus replication in nasal turbinates and lungs, but also with a reduction of virus load in the brain. Employing pseudovirus neutralization assays, we were able to identify Nbs with neutralizing capacity against the Alpha, Beta, Delta and Omicron variants. Furthermore, cocktails of different Nbs performed better than individual Nbs to neutralize two Omicron variants (B.1.529 and BA.2). Altogether, the data suggest these Nbs can potentially be used as a cocktail for intranasal treatment to prevent or treat COVID-19 encephalitis, or modified for prophylactic administration to fight this disease.

## Introduction

In 2019, Severe Acute Respiratory Syndrome Coronavirus 2 (SARS-CoV-2) emerged and was responsible for the infectious respiratory disease COVID-19, which ultimately resulted in an unprecedented pandemic that affected millions and placed a significant burden on global trade and healthcare systems. The virus spread rapidly, reaching a fatality rate between 0,4 % and 1,5% worldwide [1,2]. To date, there have been more than 759,408,703 confirmed cases and over 6,866,434 deaths, reported to the World Health Organization (WHO) [3]. In Argentina, COVID-19 reached an alarming case load with a total of 10,4 million cases and 130,463 deaths [4].

SARS-CoV-2 is a member of the *Coronaviridae* family, order Nidovirales, and genus *Betacoronavirus* [5]. The SARS-CoV-2 virion encapsidates a positive-stranded RNA genome protected by a nucleocapsid (N) and covered by a lipid bilayer envelope [6]. The envelope contains three different glycoproteins, the Spike protein (S), the Envelope protein (E), and the Membrane protein (M) [7]. The S proteins are organized as trimers on the viral envelope, with each monomer being composed of 1273 amino acids (AA) divided into two subunits: the S1 subunit, containing the receptor binding domain (RBD), and the S2 subunit that allows fusion of viral and cellular membranes. During infection, the RBD interacts with the Angiotensin Converting Enzyme 2 (ACE2) receptor located on the membrane of the target cells, with the S protein promoting viral fusion with host cell membranes to allow virus entry into the host cell [8].

Great efforts have been made to develop and test different types of vaccines to combat the ongoing pandemic. To date, there are a total of 11 vaccines commercially available [9] and 69.7% of the world population has received at least one dose of a COVID-19 vaccine [10]. However, limited immunity responses (i.e., low magnitude and short duration) to natural infection and vaccination, as well as the emergence of new SARS-CoV-2 variants of concern (VOCs), opens questions regarding the efficacy of the vaccines and highlights the need for the development of novel and more innovative prophylactic and therapeutic products [11,12].

The use of antivirals presents an excellent alternative and a complementary strategy to control the COVID-19. Several approaches using monoclonal and polyclonal antibodies or proteins, which demonstrate binding specificity and neutralizing capacities, are being explored^7,8^. Monoclonal antibodies (mAbs) that target the S or RBD proteins have been shown to be useful in treating SARS-CoV-2 infection [13,14]. However, it has been reported that the efficacy of some mAbs against specific variants and subvariants can be variable [15]. The Food and Drug Administration (FDA) has granted emergency use authorization to a number of antibody therapies, generally in the form of a mixed cocktail of antibodies, to better target a broad spectrum of SARS-CoV-2 variants [16]. One concern about employing mAbs therapies is that the emergence of new variants, such as the Omicron, are often unaffected by their use [17–19]. For this reason, other passive immunization strategies, such as the use of llama-derived nanobodies (Nbs), are an attractive alternative to develop highly sensitive diagnostic methods, and contribute to the preventive and therapeutic treatment of the disease [20].

Llama-derived Nbs are composed of only the variable region (VHHs) of the heavy chain antibodies present in the serum of camelids [21]. These antibody fragments have small molecular weight and are one of the smallest molecules known in nature to have an antigen-binding function [22]. Nanobodies have high expression yields, are easy to produce and are associated with significantly lower production cost compared to conventional mAbs [22]. More importantly, Nbs possess several unique properties compared to mAbs and derived antibody fragments, such as better stability at different pH and high temperatures, higher water solubility, and the ability to cross the blood-brain barrier (BBB) and penetrate deep into the tumors [22]. Nanobodies also have the capability to access hidden cryptic epitopes on the surface of antigens, causing viral disassembly [23]. Nanobodies are easily modifiable to improve their binding and neutralizing properties [24]. Also, due to their small size and similarities to human antibodies, they are easily humanized if needed. However, most Nbs induce low or no anti-drug antibodies (ADAs), so they have broad perspectives in the fields of pharmaceutical applications, clinical diagnostic, and therapeutics [25].

Here, we report the identification and characterization of several S protein-specific Nbs that efficiently neutralize different variants of SARS-CoV-2 *in vitro* and *in vivo*. These Nbs were obtained after creating a VHH-library derived from a llama immunized with the ectodomain of the pre-fused locked S and RBD antigens originated from the Wuhan strain of SARS-CoV-2.

## Results

### Selection of SARS-CoV-2 S-2P and RBD specific Nanobodies

Recombinant SARS-CoV-2 S-2P and RBD proteins were used as antigens for the generation and selection of specific Nbs. At the beginning of the pandemic, we received plasmids to express the S-2P protein from the Vaccine Research Center (VRC, NIH) and the Icahn School of Medicine at Mount Sinai (ISMMS) [26]. The RBD expressing vector was also kindly supplied by ISMMS. Using the HEK- 293T adherent cell line, the VRC vector coding for the S-2P protein yielded significantly higher quantities of protein than the ISMMS vector encoding a slightly different protein, 0.36 mg/l and 0.09 mg/l, respectively. RBD expression under the same conditions yielded 0.8 mg/l. Due to the low yield in the S-2P protein expression using the HEK-293T adherent system, and considering the amount of recombinant protein needed to elicit a good immune response in llamas, we decided to use another source of S protein kindly supplied by Dr. Yves Durocher, NRCC, Canada, for the last two boosters. A one-year-old male llama (*Lama glama*), seronegative for antibodies to human SARS-CoV-2, was immunized according to the schedule described (**S1A Fig**). After three immunizations, the llama developed strong antibody responses to the S-2P protein, and RBD as measured by enzyme-linked immunoassay (ELISA), with a peak IgG antibody titer of 262,144 for both antigens on post immunization day 32 (PID) (**S1B Fig**). The llama’s antibody response showed strong virus neutralizing activity, measured by virus neutralization test using pseudotyped lentiviruses (pVNT) expressing the S protein corresponding to WT SARS-CoV-2 strain (**S1C Fig**). The neutralization capacity increased after each immunization, reaching a peak at a dilution of 1:1296 from a serum sample taken on PID 32 (IC90). After confirming optimal antibody responses, the llama rested for one month to promote the hypermutation process and to improve the affinity of the humoral response. The llama received a fourth immunization dose on PID 50 and was bled four days after this final boost to produce the Nb-library. A total of 200 ml of blood was collected, yielding 4.56✕10^8^ peripheral blood lymphocytes (PBLs). RNA was extracted from 1✕10^6^ PBLs and an immune library of 1.8✕10^9^ independent transformants was obtained. All clones analyzed (48/48) had a fragment of ∼700 bp, indicating the incorporation of the coding sequence of a Nb (**S1D Fig**).

Before the pandemic started, another Nb-library, obtained after immunizing a llama with the BCoV Mebus vaccine, was available. The llama immunized with BCoV Mebus developed a strong Ab response not only to the bovine virus used for immunization, but also to the SARS-CoV-2 S-2P and RBD proteins (**S2B Fig**). However, the polyclonal Abs did not neutralize SARS-CoV-2 by pVNT (**S2B and S2C Fig**). A structural comparison of the SARS-CoV-2 S protein relative to the BCoV protein was done using Uniprot KB interprot interactive modeling. High similarities in the overall structure can be observed (**S2D-E Fig**).

Both libraries were biopanned with SARS-CoV-2 antigens to obtain the desired Nbs (**S2 Fig)**. Three consecutive rounds of panning were performed to select specific Nbs for the S-2P and RBD proteins. A total of 190 colonies were screened to assess Nb specificity for the antigenic proteins by recombinant phage ELISA (rPE) and periplasmic extract ELISA (PEE). After biopanning and screening with the S- 2P protein, 74 clones were positive for rPE and 62 for PEE. The same clones were then analyzed by ELISA using plates coated with RBD, 45 clones were positive for rPE and 27 for PEE. When the biopanning and screening were done with the RBD protein, rPE and PEE resulted in 46 and 53 positive clones, respectively. While screening with S-2P protein gave 55 and 53 positive clones for rPE and PEE, respectively. A total of 18 positive clones were obtained from the BCoV Mebus library after biopanning with S-2P and screening clones with the S-2P and RBD in rPE. Seventy-two positive clones were transformed in DH5α cells and sent for sequencing. Forty-three unique Nbs were detected from the SARS-CoV-2-library and 2 from the Mebus-library (Mebus Nb-10 and Mebus Nb-25). A sequence logo plot showing the expected variability on the complementary-determining regions (CDRs) is depicted in **S3 Fig**.

### Phylogenetic analysis, repertoire diversity and germline origin of SARS-CoV-2 Nanobodies

We constructed a phylogenetic tree with the nucleotide sequences of the 43 selected Nbs from the SARS-CoV-2 immune library using MEGA version 11 (**Fig 1**). As expected, given the large sequence variability, most of the Nbs did not form clusters. However, we identified four clusters of at least three Nbs each (**Fig 1**, A to D, highlighted in different colors). Nanobodies in all clusters were selected with both antigens, indicating a good selection strategy. Cluster A was supported with a 98% bootstrap value and constituted a monophyletic group of binders directed to RBD (Nb-27, Nb-143, Nb-106, Nb-104 and Nb-110) with 69% to 91% AA homology, all with a CDR3 of 12 AA long except for Nb-110 that has a longer CDR3 of 17 AA. Cluster B was supported with a 100% bootstrap value and included four binders directed to RBD, which possess almost the same CDR3, suggesting they belong to the clonal expansion of the same B cell. Cluster C was composed of five Nbs with similar lengths in the CDR3 and an average homology of 91%. Cluster D had a low bootstrap value of 56%, however the homology was high (82.4 to 91.2%) and all Nbs had a 19 AA long CDR3.

**Fig 1.**
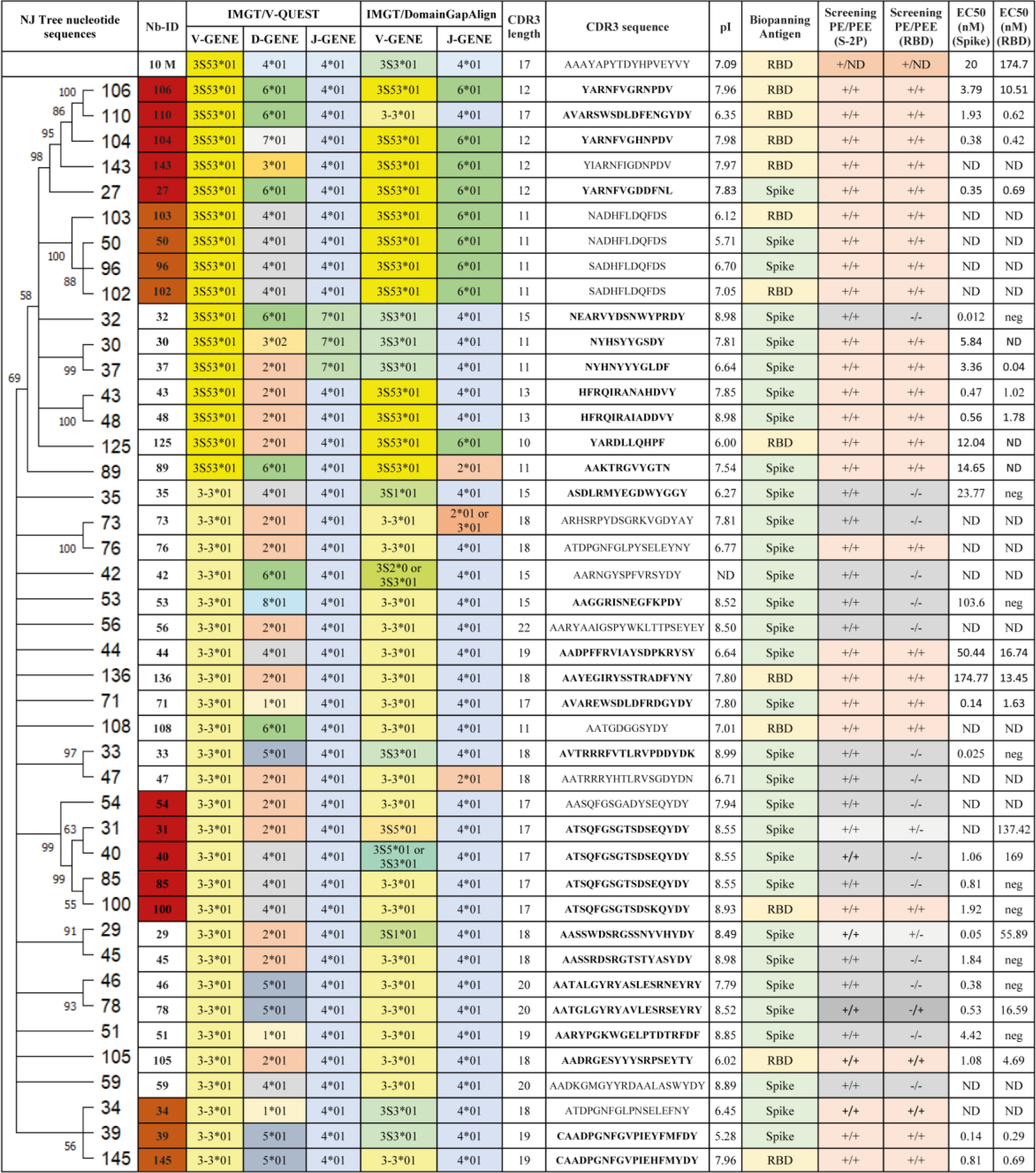
Characterization of isolated Nbs against SARS-CoV-2. Phylogenetic analysis of Nbs selected after the biopanning of two Nb-libraries, one of them generated after having immunized a llama with the bovine coronavirus (BCoV) Mebus and another with S-2P and RBD proteins of SARS-CoV-2. Nanobodies can be clustered in four groups (a, b, c and d, highlighted by the red and orange branches, column 1). Germline origin according to IMGT/V-QUEST (columns 3 to 5) and the IMGT/DomainGapAlign programs (columns 6 and 7) show gene segment (V), diversity gene segment (D), and junction gene (J) for each Nb. CDR3 composition and length are depicted in column 8 and 9. CDR3s marked in bold correspond to the Nbs selected for further characterization. Isoelectric point values are shown in column 10. The antigen used for panning and binding to the SARS-CoV-2 proteins is depicted in columns 11 to 13. The binding activity of each Nb to the S-2P and RBD proteins (half-maximal effective concentration, EC_50_ in nM), is shown in columns 14 and 15. ND: not determined.

The germline origin of the selected Nbs was analyzed by comparing their sequences with those available in two databases: IMGT/V-QUEST and the IMGT/DomainGapAlign. The IMGT/V-QUEST program, which includes only *Vicugna pacos* (alpaca) nucleotide sequences, allowed us to do a rough analysis of the V, D, and J domains’ origins. The sequences were also analyzed using IMGT/DomainGapAlign, in which the protein sequences are compared with a *Lama glama* database. In this case, only information on V and J domains can be retrieved. Results from the IMGT/V-QUEST analysis showed a predominant use of the V3S53*01 gene and allele for groups A and B, while the majority of V genes for groups C and D were V3-3*01. More variability in V gene usage was retrieved by the IMGT/DomainGapAlign analysis. When analyzing J segments, J4*01 was used in all groups according to the V-QUEST program. Nevertheless, for the DomainGapAlign program, J6*01 was predominant for clusters A and B and J4*01 for C and D. D segments have great variability according to the V-QUEST program, as can be seen in **Fig 1**. Only Nb-30 has a different allele, in this case, D3*02. The CDR3 of the selected Nbs averaged 15 (12-20) AA long and showed high sequence diversity. Considering the Nb sequences and their germinal and phylogenetic origin, we can affirm that when screening a large Nb-library (1.8✕10^9^ independent transformants) and using two selection antigens, it is possible to obtain Nbs with high sequence variability and diverse origin.

### SARS-CoV-2 Nanobody characterization *in vitro*

The selected Nbs were transformed into WK6 expression cells, then produced and purified by IMAC for further characterization. Fourteen Nbs showed notably low yield, so we selected only 29 Nbs whose level of expression ranged from 1 to 9.2 mg/l for further characterization. Eight Nbs selected with RBD recognized this domain alone or in the context of the S-2P protein. The half maximal effective concentration (EC_50_) values of these Nbs, determined by ELISA, ranged from 0.42 to 13.45 nM for RBD and 0.38 to 174.77 nM for S-2P (**Fig 1**). Twelve Nbs selected with S-2P and directed to RBD have EC_50_ values ranging from 0.04 to 137.42 nM for RBD and 0.05 to 50.44 nM for S-2P. Nine Nbs that were selected with the S-2P protein and did not bind to RBD, recognized the S-2P protein with EC_50_ values ranging from 0.025 to 103.6 nM. In particular, Nb-27, Nb-39, Nb-43, Nb-48, Nb-71, Nb-104, Nb-110 and Nb-145 had the highest affinity to the SARS-CoV-2 proteins, with the lowest EC_50_ value measured for Nb-39 of 0.14 and 0.29 nM for S-2P and RBD, respectively (**Fig 2A-B and Fig 1**). The Nbs capable of binding to S-2P in epitopes outside of the RBD, included Nb-32, Nb-33, Nb-45, Nb-46, Nb-51, and Nb-85 showed EC_50_ values below 4.5 nM **(Fig 1)**. Regarding the Nbs recovered from the BCoV Mebus library, only Nb-10 was expressed in good yield and recognized RBD alone and in the context of S-2P with EC_50_ of 174.7 and 20 nM, respectively.

**Fig 2.**
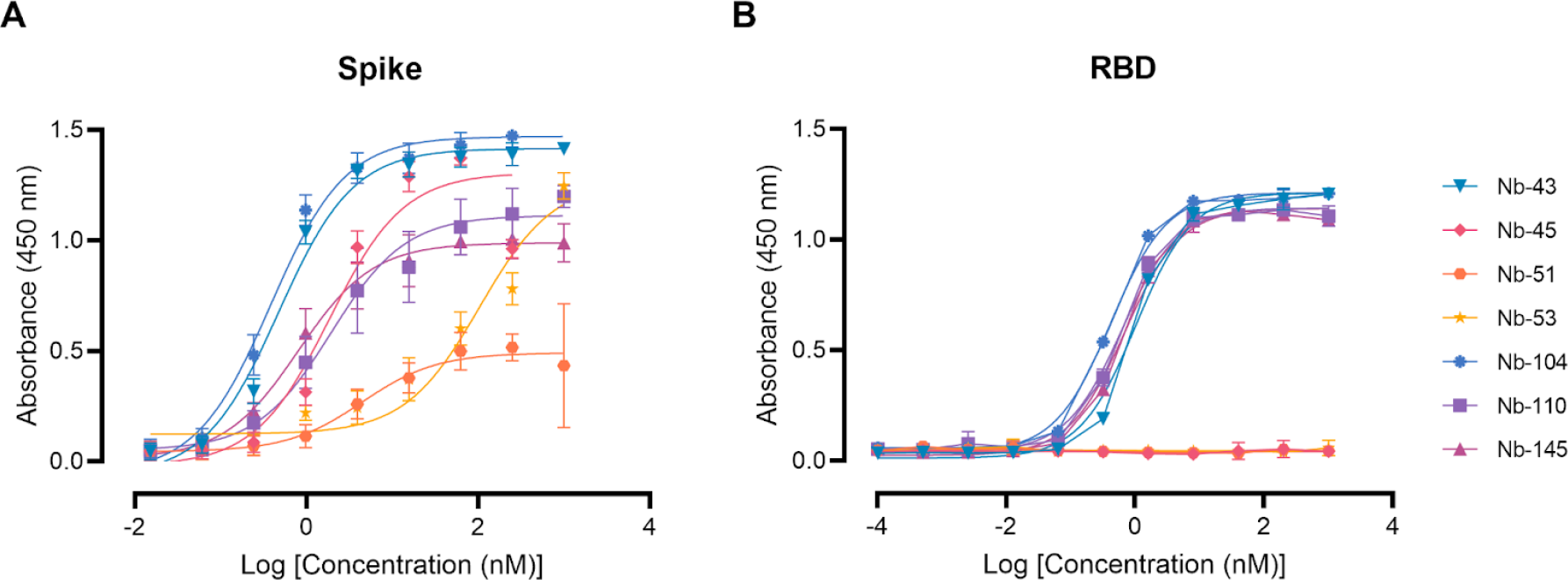
Nanobody affinity to RBD and S-2P from SARS-CoV-2 by ELISA. The binding of the selected Nbs was analyzed in plates coated with S-2P (**A**) and RBD (**B**) proteins from the WT Wuhan- Hu-1 SARS-CoV-2. Different colors were assigned to each curve according to the Nbs used. Error bars represent the standard deviation (SD) of triplicates.

### Screening of neutralizing Nanobodies against SARS-CoV-2 WT strain

A preliminary screening of the 29 selected Nbs by pVNT identified 15 potential neutralizers with half-maximal inhibitory concentrations (IC_50_) ranging from 3.36 to 79.04 nM (**Table 1**). pVNTs were performed in triplicate to determine the IC_50_. Three Nbs binding the RBD protein, Nb-104, Nb-110, and Nb-145, had IC_50_ values of 3.36, 6.05, and 24.23 nΜ, respectively (**Fig 3A)**. Five Nbs selected against the S-2P protein, Nb-39, Nb-43, Nb-45, Nb-51, and Nb-53, showed neutralization activity at 6.48, 12.08, 17.79, 54.7, and 72.68 nΜ, respectively (**Fig 3B**). The Nb-10 selected from the BCoV Mebus library, although able to recognize the RBD domain by ELISA, did not show any neutralizing properties.

**Figure 3.**
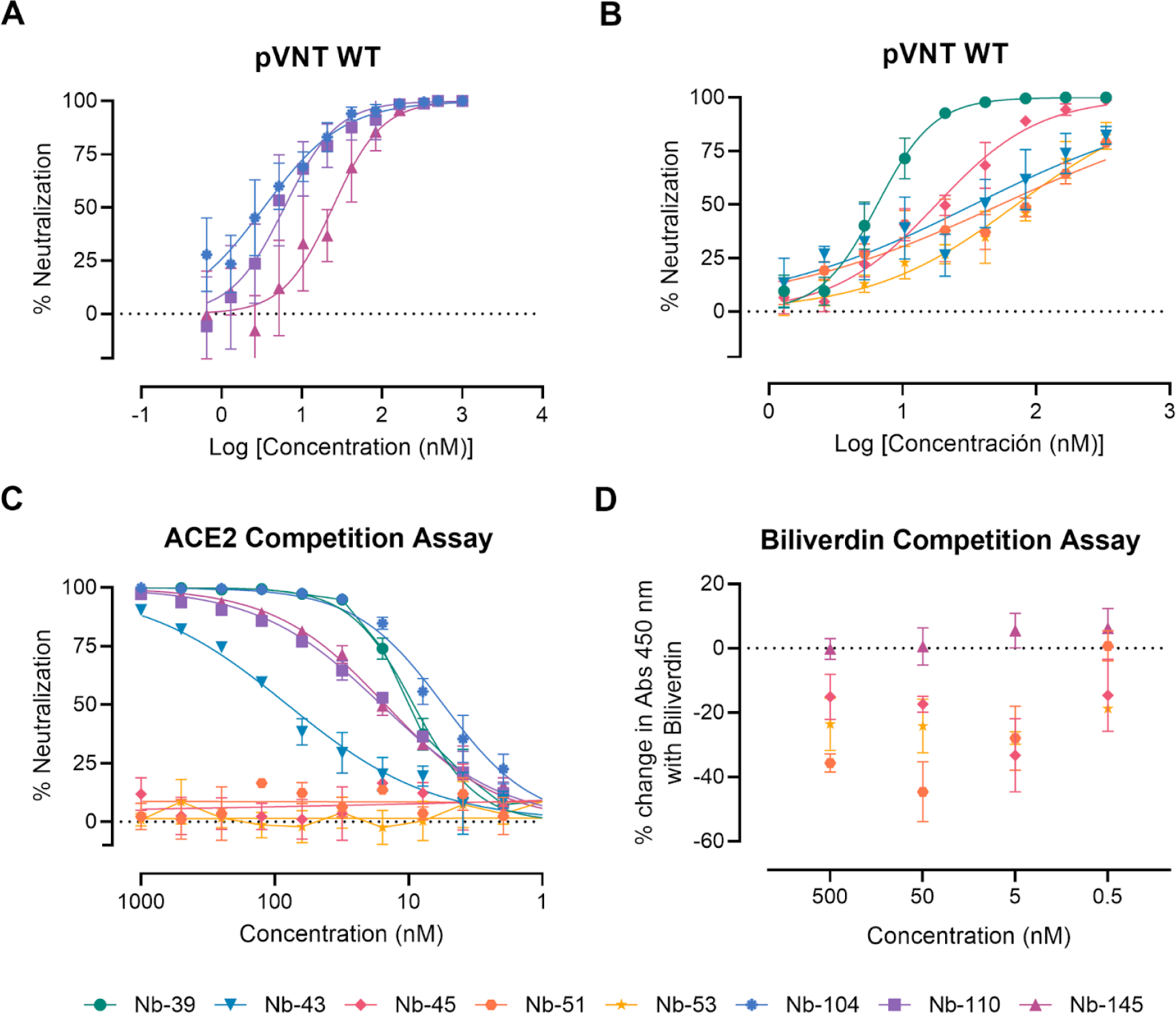
Neutralizing activity of Nbs against SARS-CoV-2 WT strain and its ability to compete with ACE2 for RBD binding or with biliverdin for S-2P interaction. Neutralization potency of eight Nbs was calculated on the basis of the pVNT results. Blue, violet, and magenta symbols and lines denote Nb-104, Nb-110 and Nb-145 directed against the SARS-CoV-2 RBD protein, respectively (A). Green, light-blue, pink, orange and yellow symbols and lines represent Nbs directed against the S-2P protein (**B**). Inhibition curves were performed with the selected Nbs at two-fold serial dilutions. After 48 h, the GFP signal from two images per well was quantified using ImageJ/Fiji and normalized to the number of GFP-positive cells of wells containing only pseudovirus. Inhibition curves are presented in log- transformed dilution with IC_50_ values for each Nb. Each experiment was replicated three times. The IC_50_ was calculated by fitting the inhibition from serially diluted Nbs to a sigmoidal dose-response curve. (**C**) Competitive ELISA of ACE2 binding to RBD immobilized on plates by increasing concentrations of Nbs. The specific binding of ACE2-HRP to RBD was detected with a chromogenic reagent. The IC_50_ was calculated by fitting the inhibition from serially diluted Nbs to a sigmoidal dose- response curve. The experiment was performed in triplicate. (**D**) Competitive ELISA of Nbs at different concentration binding to S-2P protein in the presence of 5 μM of biliverdin.

**Table 1.** IC_50_ and IC_90_ of the selected Nanobodies

| Nb-ID | IC<sub>50</sub> (nM) | IC<sub>90</sub> (nM) |  |
| --- | --- | --- | --- |
|  | pVNT | PRNT | TCID<sub>50</sub>/IF |
| 27 | 26.22 | ND | 300 |
| 29 | 74.37 | 1060 | neg |
| 30 | neg | 250 | neg |
| 31 | neg | neg | neg |
| 32 | neg | neg | neg |
| 33 | 79.04 | neg | neg |
| 35 | neg | neg | neg |
| 39 | 6.48 | 3.02 | 400 |
| 40 | neg | ND | neg |
| 43 | 12.08 | 26.9 | 100 |
| 45 | 17.79 | ND | 338 |
| 46 | 625 | ND | neg |
| 48 | 78 | ND | neg |
| 51 | 54.7 | ND | ND |
| 53 | 72.68 | ND | ND |
| 71 | 61.14 | neg | 1000 |
| 78 | 626.4 | neg | neg |
| 104 | 3.36 | 13.4 | 300 |
| 105 | 33.69 | ND | ND |
| 110 | 6.05 | 221 | 1000 |
| 125 | 67.48 | 31.25 | neg |
| 136 | 316.9 | ND | 300 |
| 145 | 24.23 | 231 | neg |

Virus neutralization activity of each Nb was assessed by three different methodologies: pseudovirus neutralization test (pVNT), plaque reduction neutralization test (PRNT) and immunofluorescence assay (IF). Half-maximal inhibitory concentrations (IC<sub>50</sub>) or ninety percent inhibitory concentration (IC<sub>90</sub>) were calculated for each Nb in each assay as described in the methods section. ND: not determined.

Neutralization ability of the selected Nbs was also determined by virus neutralization assay (VNA), applying two different methodologies, immunofluorescence (IF) and plaque reduction neutralization test (PRNT), and using two WT isolates: hCoV-19/Argentina/PAIS-C0102/2020 (containing the mutation D614G) circulating in Argentina and hCoV-19/USA-WA1/2020 from USA. The results, summarized in **Table 1**, show the neutralization capacities exhibited by 23 Nbs. As observed in the VNA, Nb-39, Nb-43, Nb-104, Nb-110 and Nb-145 exhibit significant neutralization of the WT infectious viruses. In contrast, Nb-10 selected from the BCoV Mebus library failed to neutralize both WT viruses. It is important to mention that even though a difference in the IC_50_ values can be observed for each assay, the same Nbs were identified to be neutralizing in two different tests, performed by two independent labs and using two distinct isolates.

We next examined whether the neutralizing Nbs were able to block RBD interaction with the ACE2 receptor in a competition assay, which was measured by a surrogate ELISA. Nanobody 39, Nb-43, Nb- 104, Nb-110 and Nb-145 were found to compete with ACE2 for binding to RBD with an IC_50_ of 9.16, 80.21, 5.47, 15.94, and 14.83 nM, respectively. These were classified as RBD binders. Nanobody 45, Nb-51 and Nb-53 did not compete with ACE2, suggesting that they bind to epitopes outside the RBD, and were classified as non-RBD binders **(Fig 3C**).

To further map the epitope recognized by the non-RBD binders we carried out a biliverdin competition assay, as it has been described that this metabolite binds to an epitope on NTD of the S protein and competes with a fraction of S-specific serum antibodies [27]. Our results show that the addition of 5 μM biliverdin reduced binding of Nb-45, Nb-51 and Nb-53 (at a concentration of 5 nM) to the S-2P protein by a percentage of -25.19, -20.96 and -29.24, respectively. By contrast, Nb-145 binding (RBD binder) was not affected by the addition of biliverdin (4.45%) (**Fig 3D and S4A Fig**). In a separate experiment, a dose-response assay was performed using Nbs at 100 nM and biliverdin from 12.5 to 0.1 μM. In this study, we confirmed that binding of Nb-45, Nb-51, and to a lesser extent Nb-53, was reduced in the presence of biliverdin and that this decline increased with a higher concentration of this metabolite (**S4A-C Fig**). Considering these results, we were able to predict that the neutralizing non-RBD binders might be detecting epitopes located close to the NTD of the S-2P protein.

Western Blot assays were performed to assess whether the Nbs could recognize SARS-CoV-2 S-2P and/or RBD proteins under reducing, non-reducing, denaturing or non-denaturing conditions **(S5 Fig)**. None of the selected Nbs were able to recognize RBD or S-2P proteins under reducing and denaturing conditions (**S5A Fig**). Only Nb-39 detected RBD under non-reducing and non-denaturing conditions. Non-RBD binders (Nb-45, Nb-51 and Nb-53) did not react with RBD in any of the conditions tested but they do recognize S-2P under non-reducing and non-denaturing conditions, confirming they are directed to conformational epitopes outside RBD (**S5B Fig**).

### Protection against SARS-CoV-2 challenge in k18-hACE2 mouse model

To assess the protective efficacy of Nbs against SARS-CoV-2 infection and mortality, k18-hACE2 mice were challenged intranasally with 1✕10^5^ PFU of the WT strain. According to neutralization data available at challenge, 10 or 20 μg (1 or 2 mg/Kg) of each Nb were administered to seven groups (n=9) of mice. Eighty percent of mice treated with 10 μg of Nb-39 were protected from mortality. These results were in concordance with the high VN titer of this Nb observed in the neutralizing tests conducted *in vitro*. Animals receiving 20 μg of Nb-110 and Nb-104 showed survival proportions of 60% and 50%, respectively. The group that received 10 μg of Nb-43 showed 40% survival. Only one mouse survived in the group treated with Nb-33 (non-RBD binder and non-neutralizing *in vitr*o for the challenge strain WA1/2020), while mice treated with Nb-45 (non-RBD binder, slightly neutralizing for WT strain) or irrelevant rotavirus-specific Nb-2KD1 died between days 5 and 10 post-challenge (**Fig 4B**). All mice that received the Nbs either retained or increased their body weight throughout the experiment (**Fig 4A**).

**Fig 4.**
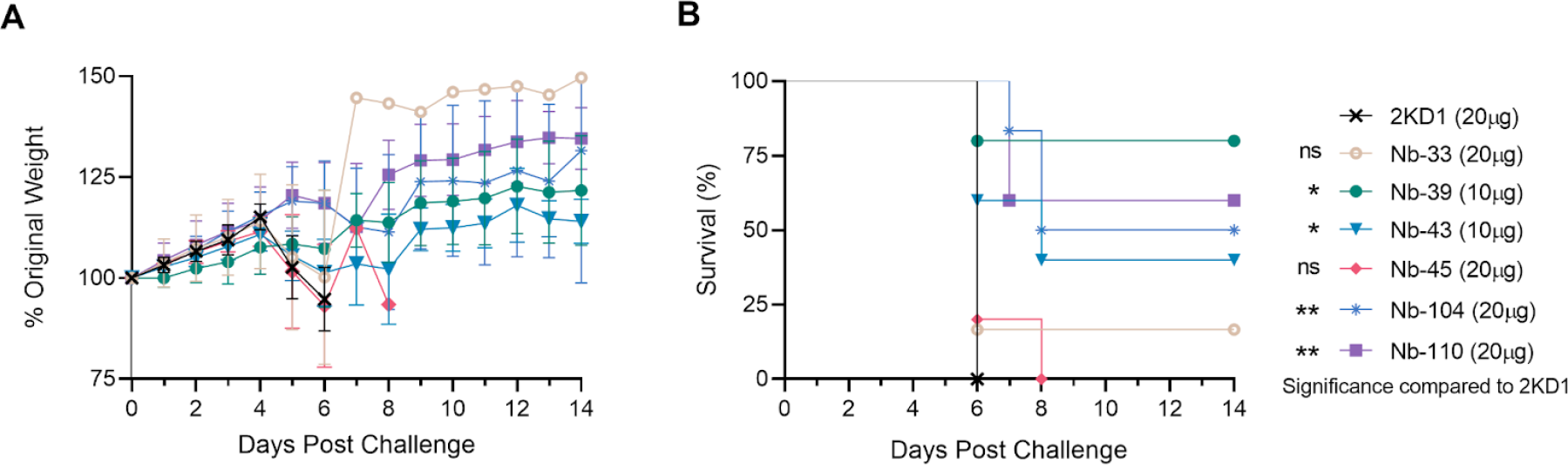
Protection against SARS-CoV-2 challenge in k18-hACE2 mouse model. k18-hACE2 mice were challenged with 1✕10^5^ PFU of with WA-1 SARS-CoV-2 after the intranasal administration of 10 or 20 μg of Nbs. (**A**) The body weight change of the animals in the control group and treatment groups was recorded for two weeks and compared. (**B**) Survival curves of the different treatment groups. Statistical significance was determined using One-way ANOVA with Dunnett’s post-hoc analysis.

Nanobody-39 and Nb-104 significantly reduced virus loads (3-4 Log_10_ titer) in nasal turbinates, lung and brain sections compared to the negative control group (rotavirus Nb 2KD1) and the non-RBD binder and non-neutralizing Nb-33 (**Fig 5A**, general mixed linear model, p<0.001). Four days post challenge (DPC), high protection (80%) in the group receiving Nb-39 was observed and was associated with a significant reduction in virus load in the nasal turbinates, lungs and brain. Nanobody 104 and Nb-110 had 50% and 40% protection, respectively, significantly reduced virus loads in the brain and lungs, and to a lesser extent in nasal turbinates. Nanobody 43 marginally reduced the virus load in all tissues. Interestingly, Nb-45 significantly reduced the virus load in the brain but not in the respiratory tract (**Fig 5**, One-way ANOVA among treatments, in each tissue, LSD Fisher, p<0.001).

**Fig 5.**
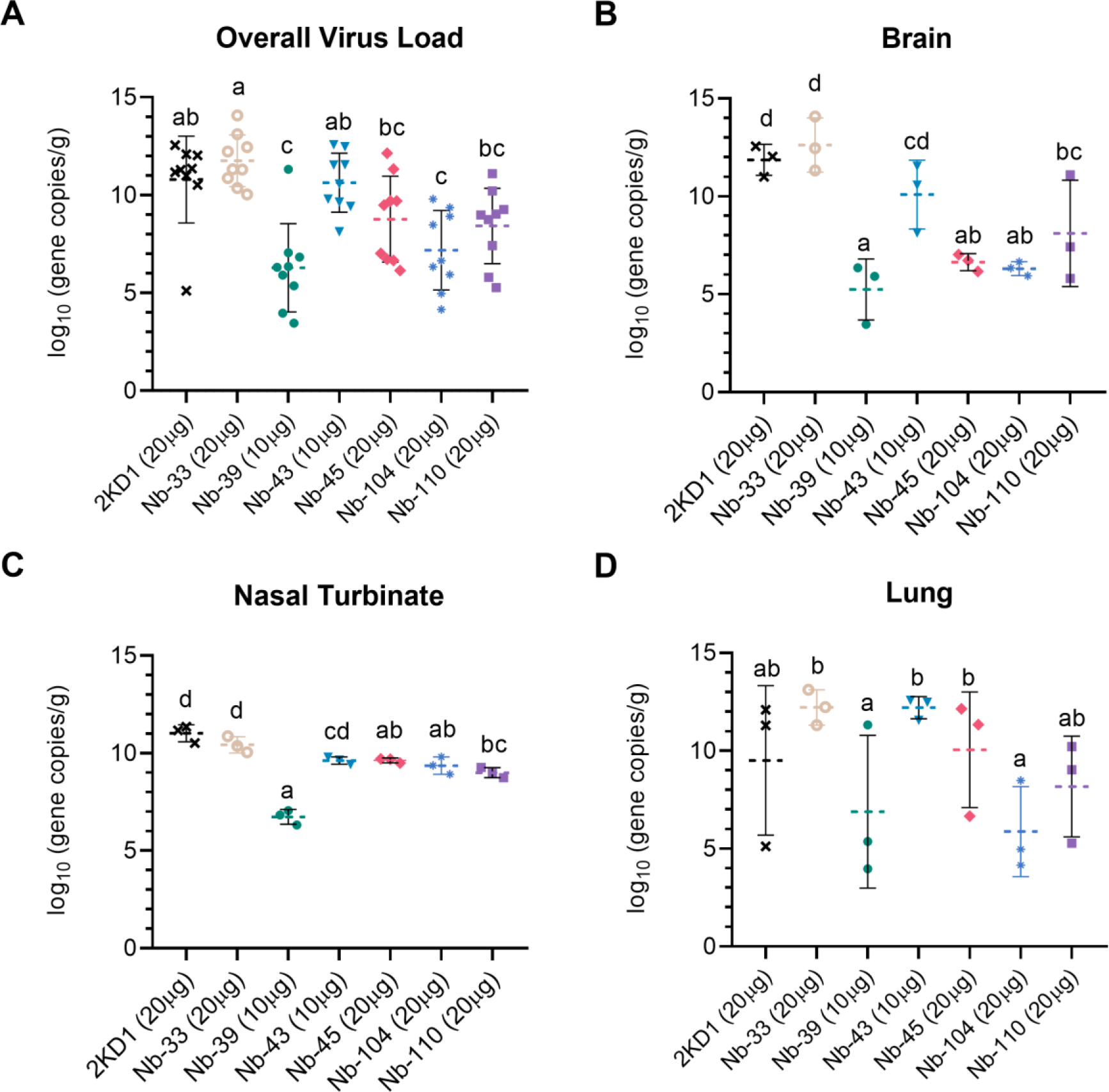
Viral load in different tissues after Nanobody treatment and virus challenge. Overall mean virus load (upper and lower respiratory tract and brain) (**A**) or virus load measured by RT qPCR in the brain (**B**), nasal turbinates (**C**) and lung (**D**) tissues collected at four days after challenge (n=3 mice per tissue, 9 mice per group). Mice were treated with each Nb by the intranasal route and challenged with WA-1 SARS-CoV-2, 4 h later. Samples were prepared from infected mice for RNA isolation and RT-qPCR. Statistical significance was determined using the general mixed 2-way ANOVA model, LSD Fisher’s exact test and Bonferroni correction. Treatment groups in the same tissue with different letters differ significantly (p<0.001).

### Nanobodies to SARS-CoV-2 WT strain neutralize SARS-CoV-2 variants

We next studied the neutralizing breadth of RBD- and non-RBD binders against SARS-CoV-2 variants. Pseudoviruses expressing the S protein of the Alpha (B.1.1.7), Beta (B.1.351), Delta (B.1.617.2), and Omicron (B.1.1.529 and BA.2) variants were prepared. Neutralizing capacity of Nb-43 (RBD-binder) was observed only for the Omicron variants B.1.1.529 and BA.2, with IC_50_ of 394.7 and 204.8 nM, respectively. Nanobody 104 (RBD binder) was able to neutralize Alpha (IC_50_=18.19 nM) and Delta (IC_50_=121.8 nM), while Nb-110 and Nb-145 (RBD binders) neutralized Alpha, Beta, and Delta variants with IC_50_ ranging from 86.65 to 451.6 nM (**Fig 6**). Interestingly, none of the non-RBD binders were able to neutralize the Delta variant. Nanobody 51 only neutralized the Alpha variant (215.5 nM), while Nb-45 and Nb-53 were able to neutralize Alpha, Beta and Omicron variants with IC_50_ ranging from 83.66 to 235.7 nM (**Fig 7**).

**Fig 6.**
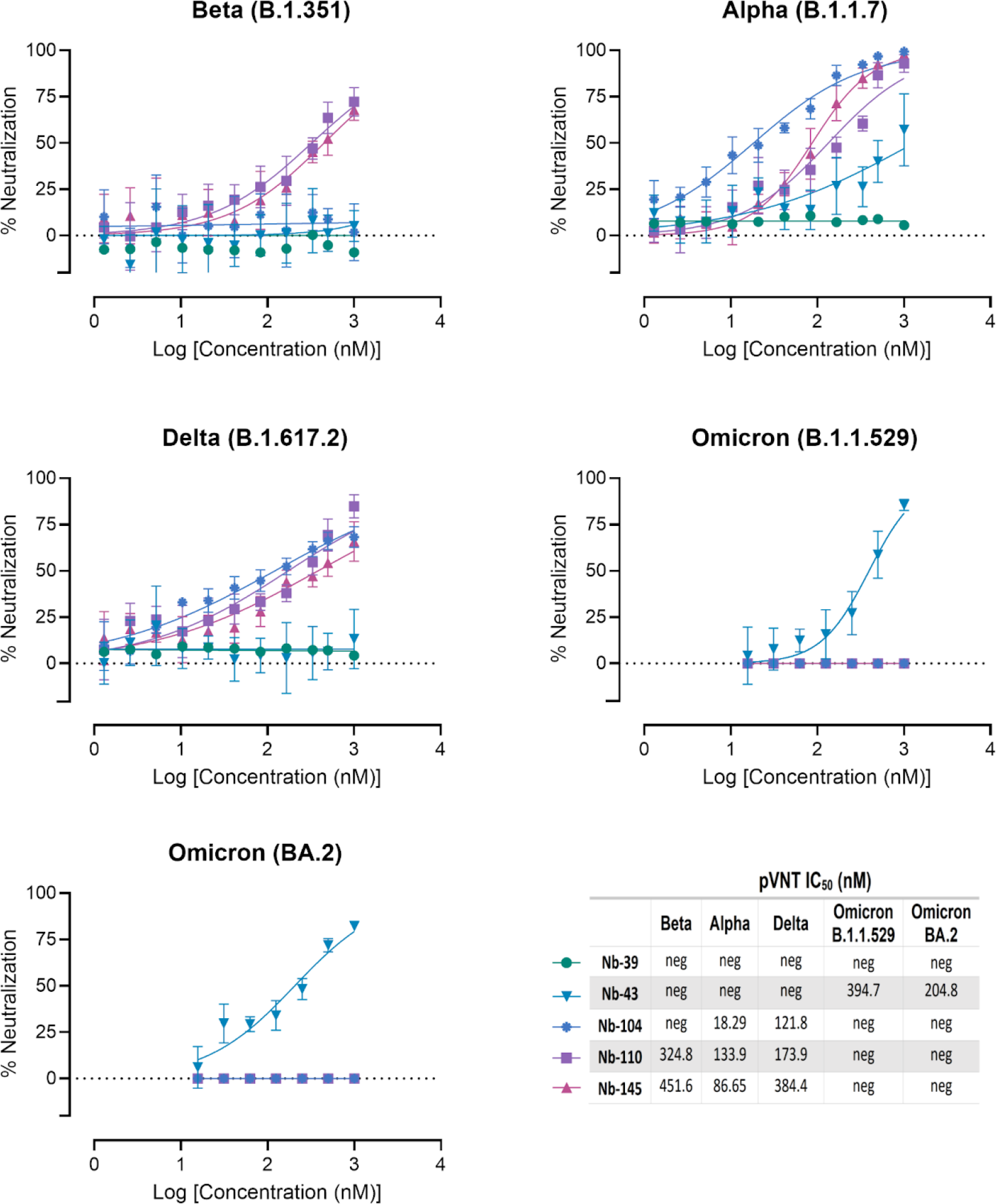
Neutralization titers of SARS-CoV-2 strains *in vitro* by RBD binders. Neutralization curves and IC_50_ of the RBD binders for Beta (B.1.351), Alpha (B.1.1.7), Delta (B.1.617.2) and Omicron (B.1.1.529 and BA.2) variants of SARS-CoV-2 measured by pVNT. Data are shown as the mean (n = 3) with SD. The IC_50_ was calculated by fitting the inhibition from serially diluted Nbs to a sigmoidal dose-response curve. The experiment was performed in triplicate.

**Fig 7.**
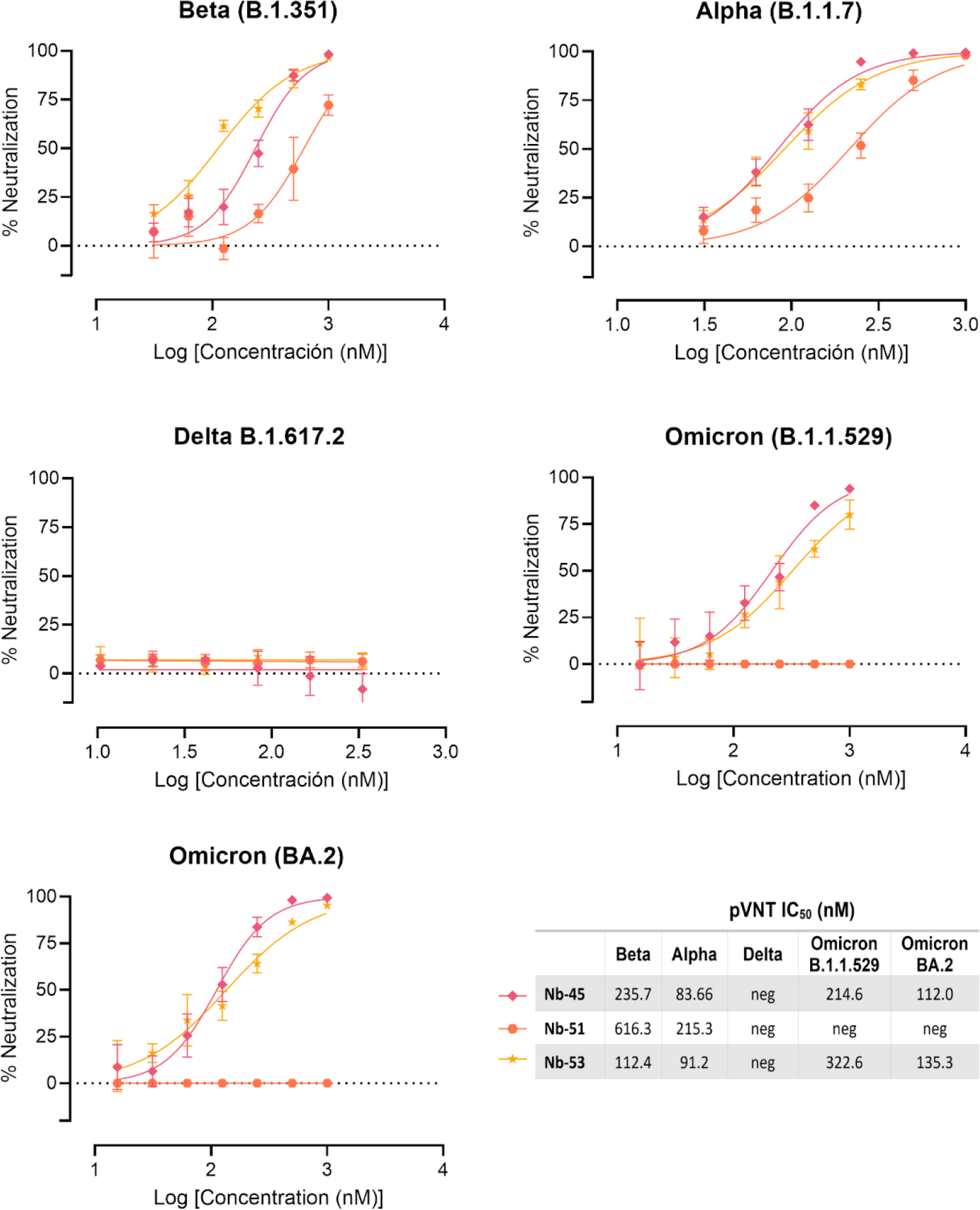
Neutralization titers of SARS-CoV-2 strains *in vitro* by non-RBD binders. Neutralization curves and IC_50_ of the non-RBD binders for Beta (B.1.351), Alpha (B.1.1.7), Delta (B.1.617.2) and Omicron (B.1.1.529 and BA.2) strains of SARS-CoV-2 measured by pVNT. Data are shown as the mean (n = 3) with SD. The IC_50_ was calculated by fitting the inhibition from serially diluted Nbs to a sigmoidal dose-response curve. The experiment was performed in triplicate.

### A cocktail of Nanobodies enhances neutralization potencies against Omicron variants

Finally, we decided to focus on the Nbs that were able to neutralize the Omicron variants, as these variants were recently circulating in our population. We tested a cocktail of Nb-43 (RBD binder), Nb- 45 and Nb-53 (non-RBD binders). When using the B.1.1.529 Omicron variant, Nb-43 did not modify its neutralizing capacity when combined with Nb-45 (394.7 to 348.4 nM), but significantly increased its neutralizing capacity to 126.8 nM together with Nb-53 (One-way ANOVA, Tukey HSD, p<0.001). Nanobody 45 alone has an IC_50_ of 214.6 nM and decreased to 171.8 nM when mixed with Nb-53, however this reduction was not significant. When Nb-45, Nb-51 and Nb-53 were mixed, a significant increase in their neutralizing capacity was observed compared with Nb-43 or Nb-53 alone, p<0.001. We did not observe an enhancement in the neutralizing capacity of the three-Nb mixture compared with the two-Nb mixtures, except for the combination of Nb-43+Nb-45, p<0.001 (**Fig 8A and C**). For the BA.2 Omicron variant, Nb-43 significantly reduced its IC_50_ value from 204.8 to 54.39 nM when combined with Nb-45 (p<0.001), and to 81.4 nM when combined with Nb-53 (p<0.001). Nanobody 45 did not improve its neutralizing capacity when combined with Nb-53. The neutralization potency increased significantly when Nb-45, Nb-51 and Nb-53 were added together compared with each Nb alone (p<0.001). In contrast to the results obtained for the B.1.1.529 Omicron variant, the three-Nb mixture, compared with the two-Nb mixture, showed a significant difference in its IC_50_ value for the BA.2 Omicron variant, especially for the Nb-43+Nb-53 and Nb-45+Nb-53 mixtures, p<0.001 (**Fig 8B and D**).

**Fig 8.**
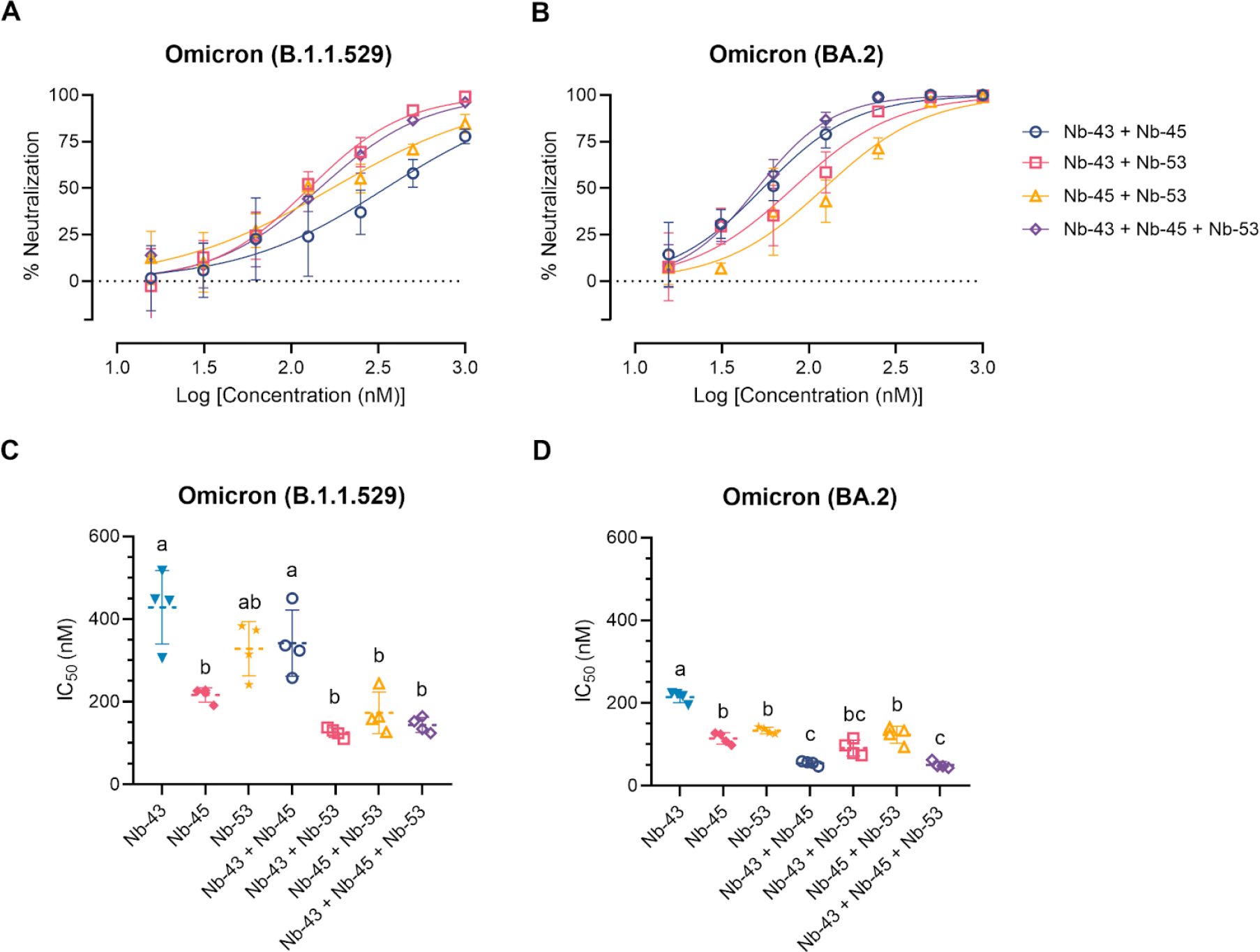
**Enhancement of neutralization potencies against Omicron variants using Nanobody cocktails**. (**A**) Percentage of neutralization exerted by a mixture of two or three Nbs against the B.1.1.529 Omicron variant. (**B**) Similar experiments than in A performed with the BA.2 Omicron variant. Comparison of IC_50_ values of Nbs combinations vs. individual Nbs for the B.1.1.529 Omicron variant (**C**) and the BA.2 Omicron variant (**D**). Individual Nbs or Nb-mixtures with different letters differ significantly (One-way ANOVA, Tukey multiple comparison, p<0.001).

### Nanobody epitope mapping and binding mode by *in silico* **prediction**

We tried to identify the target epitopes of each set of Nbs using a peptide array covering the entire AA sequence of the S glycoprotein of the USA-WA1/2020 strain of SARS-CoV-2 (BEI resources NR- 52402). Peptides were 17- or 13-mers, with 10 AA overlap. Most of the Nbs failed to react with the peptides, except for Nb-39 and Nb-33. Nanobody 39 recognized a peptide between AA 407 to 423 corresponding to the receptor binding motif (RBM) inside RBD. Nanobody 33 bound to one peptide between HR1 and HR2 region, both within the S2 domain, in agreement with the result that this Nb was selected with the S protein and did not react with RBD in ELISA (**Table 2**).

**Table 2.** Identification of target epitopes in the S protein by a peptide array

| Nanobody | Peptide | Amino acid sequence | OD 405 nM | Location |
| --- | --- | --- | --- | --- |
| **Nb-33** | 161 | 1121-FVSGNCDVVIGIVNNTV-1137 | 0.719 | S2 |
| **Nb-39** | 59 | 407-VRQIAPGQTGKIADYNY-423 | 0.423 | RBM |

Epitope mapping using a 181-peptide array spanning the S protein of the USA-WA1/2020 strain (GenPept: QHO60594). Responsive peptides were detected only for Nb-33 in the S2 domain (161) and Nb-39 in the RBM (59).

According to ELISA, WB and biliverdin competition assay results, Nb-43 is predicted to bind to the RBD, while Nb-45 and Nb-53 recognize the NDT of the S protein. A combination of these Nbs in a cocktail improved their neutralizing capacity against the Omicron variants. Considering these results, all three Nbs were selected and modeled for binding mode prediction using High Ambiguity Driven protein-protein Docking (HADDOCK) methodology.

HADDOCK provided multiple results with varying scores, we selected the best pose which positioned the Nbs’ CDRs in direct contact with the S protein (**Fig 9 and S1 Table)**. Our analysis of the interacting residues between Nb-43 and the S protein suggested that the binding site would be located at the binding interface of the complex. Residues 469, 511 and 512 of the RBD, together with residues 29 (CDR1), 104 and 105 (CDR3) of the Nb-43, may contribute as critical residues for binding affinity (**Fig 9A**). Other residues, 463 to 511, within a distance less than 5 Å might also be interacting with the Nb residues. For the non-RBD binders (Nb-45 and Nb-53), the analysis of the predicted interactions allowed us to identify potential interacting residues in the NTD of the S protein. Residues 54 (CDR2), 101 and 114 (CDR3) of Nb-45 may be interacting with residues at positions 60, 244, 249, respectively (**Fig 9B**). In case of the Nb-53, possible residues involved in the interaction Nb-NTD might be 54 (CDR2), 106 and 107 (CDR3) and 116, 206, 231, respectively (**Fig 9C**). Other NTD residues located less than 5 Å are 62, 191, and 245 to 249 for Nb-45 and 115, 117, 205, 227 to 230 and 236 for Nb-53. We also performed a HADDOCK analysis of biliverdin and observed that the predicted binding site of Nb-45 and of this molecule overlaps. When similar analysis was done for Nb-53, we observed a partial overlap with the biliverdin recognition site. Our findings support the competition assay results (**Fig 2D**) but further mutagenesis analysis and crystallographic studies are needed to confirm the residues involved in the S- Nb interaction.

**Fig 9.**
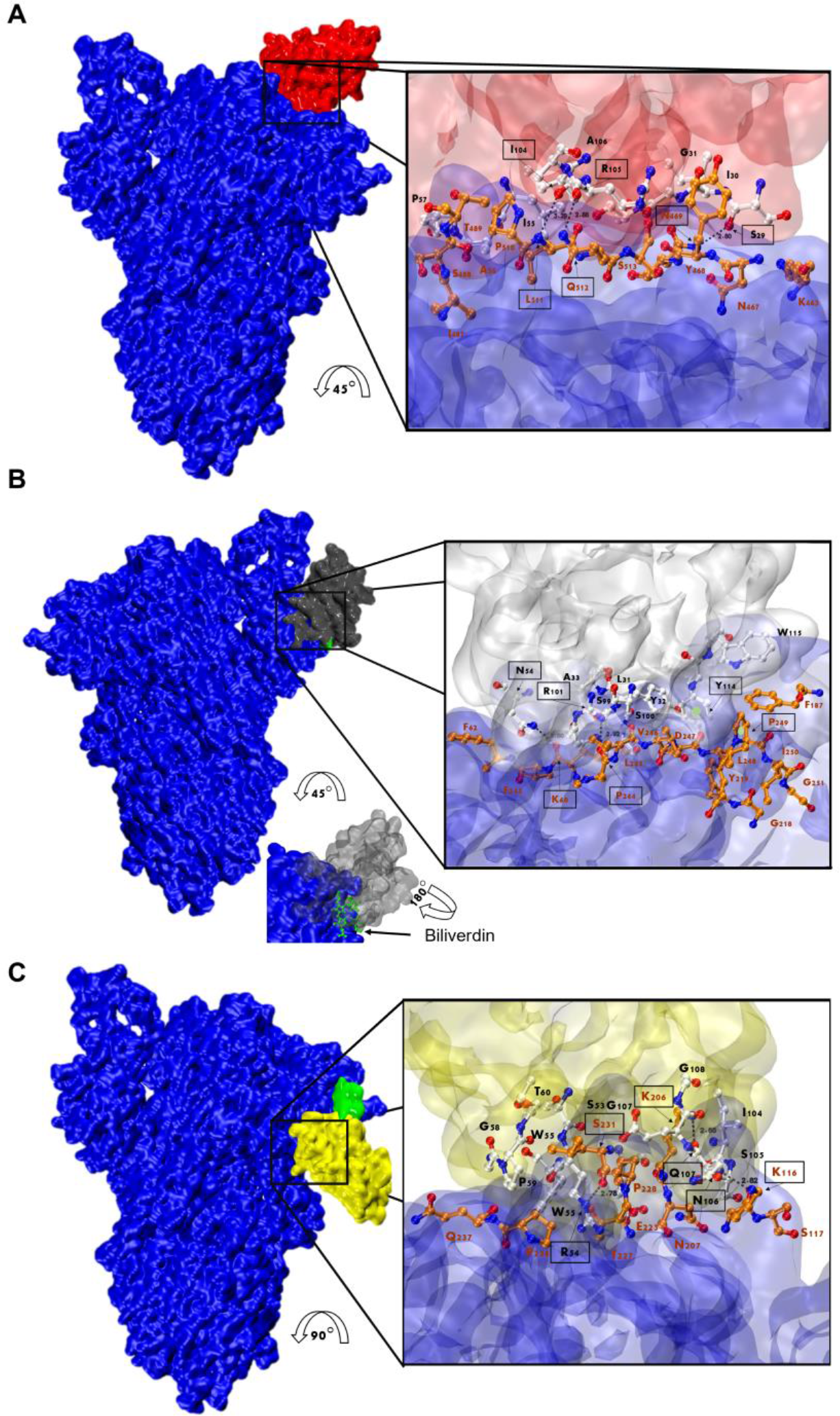
Analysis of Nanobody-S protein interactions. Prediction of the interaction of Nb-43 (**A**, red), Nb-45 (**B**, gray) and Nb-53 (**C**, yellow) with the S protein (blue). Biliverdin is highlighted in green. The three panels display the S-Nb overall view and a close-up view of the interaction zone. The close-up view displays the predicted binding site, with AA in red representing those of the S protein and in black those belonging to the Nb. Amino acids enclosed in a black box and indicated with an arrow shows residues that could form hydrogen bonds (dotted lines). Section B shows green circles representing possible Pi-stacking interactions.

## Discussion

After the onset of the COVID-19 pandemic, research groups working with camelid-derived antibodies aimed to develop Nbs to combat SARS-CoV-2 infection. This was achieved through biopanning of existing libraries for MERS and SARS coronavirus or immunizing llamas and alpacas with multiple SARS-CoV-2 antigens, primarily S and RBD recombinant proteins [28–37] . So far, several Nbs have been developed with promising diagnostic and therapeutic applications [20,38]. Both monomeric Nbs and engineered molecules showed strong neutralizing activity to the different VOCs that have emerged, including the Omicron variants [39–46]. The intense production of new recombinant Nbs highlights the relevance of this technological platform, which is expected to reach clinical use for SARS-CoV-2 and other viral diseases in the near future [47]. Here, we report the development and characterization of a novel and diverse set of unique Nbs (pending patent WO 2022/140422) with strong neutralizing activities against SARS-CoV-2 variants. Some of them were even able to neutralize the Omicron variants when combined in a cocktail or to significantly reduce viral loads in the brain.

Initially, we aimed to isolate Nbs against SARS-CoV-2 S-2P and RBD recombinant proteins from a preexisting VHH-library produced against BCoV Mebus strain, as the S protein from both strains shows high similarities in the overall structure (**S2D Fig**). Interestingly, we were able to retrieve a Nb (Mebus Nb-10), which binds to the RBD of SARS-CoV-2 alone and within the context of the S-2P protein. However, Nb-10 did not show neutralizing activity against any of the SARS-CoV-2 variants tested. Thereafter, a novel set of Nbs was developed from an immune library of a llama immunized with three types of full-length S-2P protein and recombinant RBD, all derived from the original SARS-CoV-2 Wuhan-Hu-1 strain.

After a four injections immunization protocol, an exceptionally large immune library of 1.8✕10^9^ individual transformants was generated. We obtained different groups of Nbs that exhibited high sequence variability and were originated from a diverse set of V, D, and J genes (**Fig 1**). Our results are consistent with those obtained by other groups screening camelid-derived libraries, where an impressive diversity in the Nb nucleotide sequence, specificity and neutralizing capacity was found, especially after successive immunizations to promote the development of superimmunity [48,49]. This is particularly relevant considering the emergence of new VOCs, since the screening of a large library, like the one we described in this work, constructed from hyperimmunized animals, can broaden the type of Nbs obtained and increase the probability of finding binders that recognize new variants.

We characterized 29 out of 43 unique Nbs to determine their binding and neutralizing properties. Network analysis showed that the Nbs are arranged into several clusters sharing different properties, such as target recognition, neutralization potency, CDR3 length and isoelectric point (pI) (**Fig 1 and 10**). It is worth highlighting that we isolated Nbs with similar sequence identity using two different antigens for the biopanning. Three of the largest clusters are composed of Nbs with strong neutralizing activity (IC_50_ below 25 nM) and wide breadth of neutralization of SARS-CoV-2 variants. Most of the neutralizing Nbs selected from the SARS-CoV-2 immune library are directed to conformational epitopes in the RBD domain, as shown by Western blot (**S5 Fig**). Several Nbs cannot be clustered due to their unique CDR3 sequence. For instance, Nb-53, a non-RBD binder, was able to neutralize almost all the SARS-CoV-2 variants tested, even though it had an intermediate IC_50_ value. Nanobodies with neutralizing properties that do not bind to RBD (Nb-45, Nb-51, and Nb-53), recognize only S-2P by WB and do not block the interaction with ACE2 (**Fig 2 and 3B-C**) were further studied to evaluate if their binding site was located in the NTD region. For this, we performed a biliverdin competition assay and confirmed that Nb-45, Nb-51, and Nb-53 did compete with this molecule to different degrees, supporting the hypothesis of their binding to the NTD. Neutralizing antibodies directed to this domain have also been reported by others [27,50]. Even though crystallographic and structural experiments would be needed to fully understand the Nb-NTD interaction, epitope binding prediction suggested that Nb-45 and Nb-53 are indeed interacting with this domain (**Fig 9B and C).**

**Fig 10.**
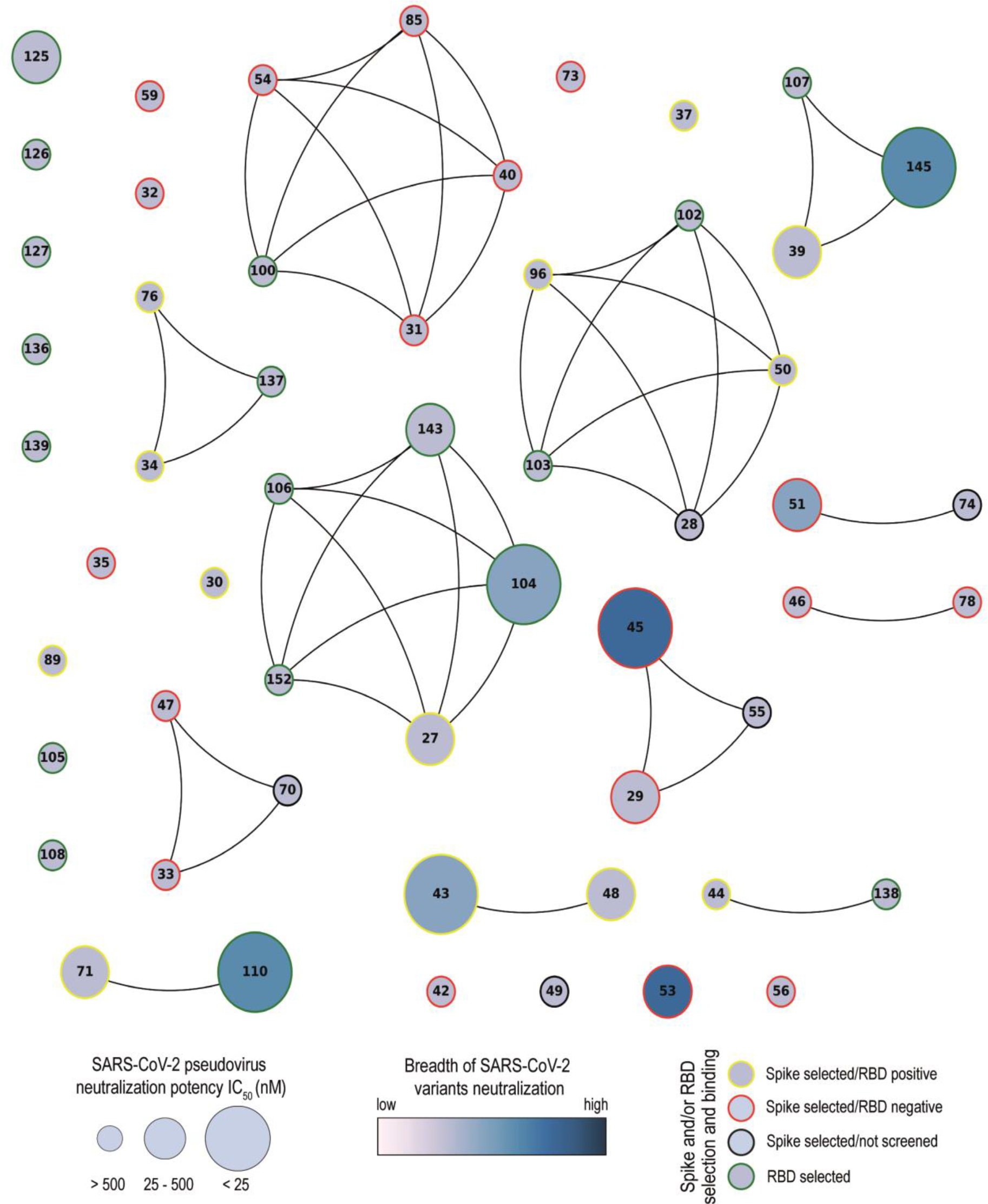
Network analysis of unique Nanobodies. A visual representation summarizing the RBD and S-2P binding and neutralization for the 43 isolated Nbs. Nanobodies are depicted as dots in different sizes and colors, and those with a CDR3 sequence identity greater than 70% are connected. Their neutralization potencies against pseudotyped SARS-CoV-2 WT are represented by the size of dots and the filled gradient color represents the breadth of SARS-CoV-2 variants neutralization. Dots are colored on the outer circle based on the antigen used for biopanning and whether they bind to RBD or not.

Nanobodies often recognize conformational epitopes, although some exceptions to this have been reported and few Nbs were able to recognize linear peptides [51,52]. In this regard, Nb-39 recognized a linear peptide corresponding to the RBM inside the RBD in an S peptide-array. This result agrees with the fact that this Nb was able to block the interaction between RBD and ACE2 together with its neutralizing activity *in vitro* and protection in mice *in vivo*. As expected, Nb-33 (non-RBD binder), recognized a peptide between HR1 and HR2 region, within the S2 domain. These regions within S2 are highly conserved and immunogenic, playing an important role in the fusion process before internalization by endocytosis, even more, human mAbs directed to these regions are usually neutralizing [53,54]. Nanobody 33 was marginally neutralizing when measured by pVNT but did not neutralize the WT virus. Furthermore, Nb-33 is especially suitable for developing diagnostic assays like biosensors for virus detection in wastewater, food and clinical samples, as it was shown that this Nb detected all S variants in a diagnostic assay (He *et al.,* submitted March 2022). Overall, these results suggest that our Nbs recognize not only conformational epitopes on the surface of the S protein but they also interact with linear regions of the protein.

Nanobodies have shown promise as prophylactic or therapeutic tools against SARS-CoV-2 infection, given their high specificity and low immunogenicity [47]. In this study, we evaluated the efficacy of Nb-33, Nb-39, Nb-43, Nb-45, Nb-104 and Nb-110 in mice as an intranasal prophylactic treatment. The RBD binder Nb-39 showed high neutralizing activity *in vitro* and gave the highest protection rate with 80% survival in mice, even though a dose of only 10 μg was administered. Nanobody 43, Nb-104 and Nb-110 showed different degrees of protection *in vivo,* while animals receiving the non-RBD binders, Nb-33 and Nb-45, did not survive. This protection was associated with a reduction of virus load in both the upper and lower respiratory tract, as well as in the brain (**Fig 5A**). In this regard, it has been reported that Nbs can cross the BBB after some modifications [55]. Furthermore, a study by Pierre Lafaye’s team has revealed that Nbs with a high isoelectric point (pI∼9.0) spontaneously cross the BBB [56]. Such Nbs not only gained access to the brain but were even found to penetrate cells and bind to intracellular proteins. Nanobody 45 and Nb-104 have the highest pI (8.98 and 7.98, respectively), and their potential capability to cross the BBB could explain the reduction of virus load in the brain, even when Nb-45 did not show a significant decrease of viral titer in the lung. These results suggest that although the neutralization exerted by Nbs in other tissues could lead to an overall lower viral load, neutralization directly in the brain cannot be ruled out. The capability of Nbs to cross the BBB could be of high impact when encephalitis has been diagnosed in patients with acute and long-term COVID [57,58]. In contrast to this hypothesis, Nb-39 showed an acidic pI (pI: 5.28) and was also able to reduce the virus load in the brain. In this case, the high reduction of virus replication in the respiratory tract might reduce virus dissemination to other organs, including the brain. Although our hypothesis regarding Nbs access to the brain needs to be confirmed by radiolabeling biodistribution assays, our data suggest that intranasal administration of a cocktail of Nbs could be used to prevent or treat COVID-19 encephalitis. This statement is further supported by other studies that have reported that Nbs can reach the brain after intranasal administration [59–61]. To our knowledge, this is the first report of SARS-CoV-2 replication reduction in the brain after a Nb treatment in a mouse model.

The emergence and spread of SARS-CoV-2 variants that can escape the neutralization by Abs pose a challenge for the development of effective vaccines and therapeutics [14]. For this reason, Nb cocktails, composed of two or more Nbs that recognize different epitopes on the S protein, have been suggested as a strategy to increase the neutralization potency and prevent viral escape [62]. We determined the neutralizing potential of Nbs in a cocktail against Omicron variants. A mixture of a RBD binder, Nb- 43, and two non-RBD binders, Nb-45 and Nb-53 were tested. Our study shows that most combinations of Nbs performed better when added in pairs rather than as individual Nbs to neutralize both Omicron variant. Addition of a third Nb improved the neutralizing capacity against Omicron B.1.1.529 only when compared with the two-Nb mixture Nb-43+Nb-45. While the three-Nb mixture performed better than the two-Nb mixture Nb-43+Nb-53 and Nb-45+Nb-53 for the BA.2 Omicron variant. Zhao and coworkers have reported the development of a Nb capable of binding to both RBD and NTD of the S trimer through the same CDR3 loop, with broadly neutralizing activity [63]. These findings highlight the potential of Nbs cocktails as a strategy for developing novel prophylactic and therapeutic tools against SARS-CoV-2, particularly in the context of emerging viral variants.

The Nbs developed in the present work react with a broad range of different epitopes that support their use in a cocktail or as multivalent engineered molecules to detect and control the SARS-CoV-2 disease. More importantly, the selected Nbs possess strong neutralization potency *in vitro* and *in vivo* in their monomeric form. However, most anti-SARS-CoV-2 Nbs that have been developed by others were multimerized to improve the potency of their neutralizing activity through an increased avidity effect [29,33,35,39,64–66]. For example, gain in KD value, increased affinity and more potent neutralizing capability has been reported for Nb trimerization compared with dimerization [36,66]. We hope to be able to test if multivalent Nbs can result in molecules with better and broader inhibitory capacity. This is feasible as it has been reported that the surface of the RBD is large enough to accommodate several Nbs at a time [67]. Therefore, Nbs that bind to different epitopes, have strong neutralizing properties and are able to reduce the viral load in the brain represent promising tools for developing a therapeutic formulation to neutralize new emerging VOCs, prevent viral escape and possibly treat COVID-19 encephalitis.

## Materials and methods

### Cells and virus

HEK-293T cells were grown in high glucose Dulbecco’s modified Eagle’s medium (Thermo Fisher Scientific, 12100046) supplemented with 10% fetal bovine serum (Natocor), penicillin/streptomycin (Thermo Fisher Scientific, 10378016) and 110 mg/l of sodium pyruvate (Thermo Fisher Scientific, 11840014). Coronavirus Isolate hCoV-19/USA-WA1/2020 (BEI, NR-52281) and Argentinian isolate hCoV-19/Argentina/PAIS-C0102/2020 (D614G), adapted to grow in Vero cell line, were used for Virus Neutralization Assay (VNA). Isolate hCoV-19/USA-WA1/2020 was also used for mice challenge.

### Recombinant protein expression and purification

Mammalian expression plasmids encoding for SARS-CoV-2 S-2P protein containing a C-terminal 6xHis tag were kindly provided by Dr. Karin Bok (VRC7473-2019 nCoV S-dFurin-WT-F3CH2S_JSM) and Dr. Florian Krammer (pCAGGs nCoV19). RBD expressing vector (pCAGGs nCoV19, residues 319 to 541) was kindly provided by Dr. Florian Krammer [26]. Plasmids were transiently transfected in HEK-293T cells using polyethyleneimine (PEI, PolyAR) in a 1:2.5 DNA:PEI ratio in OptiMem medium (Thermo Fisher Scientific, 22600050). The supernatant containing recombinant proteins was harvested 72 h post-transfection and centrifuged at 10,000 g for 20 min at 4°C, and then filtered through a 0.22 μm membrane and incubated with nickel affinity resin (Amintra, ANN0025). Proteins were purified by gravity flow using Poly-Prep® Chromatography Columns (Bio Rad, 7311550) and eluted using 300 mM Imidazole. Fractions containing the recombinant proteins, as judged by the SDS-PAGE analysis, were pooled, dialyzed against phosphate buffer saline (PBS) and stored at -80°C. SARS-CoV- 2 S-2P protein expressed in a CHO-pool system was produced in Dr. Yves Durocher at the National

Research Council of Canada as it has been previously described [68]. The expression of RBD in yeast (yRBD) was done by the Argentinian consortium using the X-33 *P. pastoris* strain as it has been previously described [69].

ACE2s-HRP was cloned in the pCAGGs vector under the β-actin/rabbit β-globin promoter. A human serum albumin secretion signal and a 6xhis tag were added at 5’ and 3’, respectively. The ACE2 sequence was cloned from ACE2_TM_3CHS vector kindly provided by the VRC using the following primers: Fw ACE2BspEI 5’-aaaTCCGGACAGTCCACCATTGAGGAAC-3’ and Rv ACE2EcoRV: 5’-tttGATATCccGCTTTGGTCTGCATATGGAC-3’. The HRP coding sequence, codon-optimized for mammalian cell expression, was added at the C-terminal on the *Eco*RV site. After searching clones with a correct orientation by restriction analysis, plasmids were amplified and transfected on HEK-293T cells and the ACE2-HRP protein was produced and purified as it was described for the S-2P protein.

### Llama immunization and library construction

A llama located at INTÁs Camelids Experimental Unit was intramuscularly immunized with 200 μg per dose of recombinant SARS-CoV-2 S-2P protein produced in HEK-293T cells on days 0 and 14 and 200 μg of SARS-CoV-2 S-2P produced in CHO cells and 100 μg of RBD protein produced in HEK- 293T cells on days 28 and 50. Two different sources of S-2P protein were used due to the low performance obtained in adherent cells (HEK-293T) and the urgency in protein availability for the immunizations. Complete Freund’s adjuvant was used for priming and Incomplete Freund’s adjuvant for the following boosters. The antibody responses to S-2P and RBD were monitored by ELISA and pVNT on day 4 and 7 post each inoculation.

Llama management, inoculation, and sample collection were conducted by trained personnel under the supervision of a Doctor in Veterinary Medicine in accordance with Argentinean and international guidelines for animal welfare. This study was approved by the Animal Care and Use Committee of INTA (CICUAE) under protocol N° 15/2020.

Four days after the last boost, 200 ml of blood was collected, and peripheral blood lymphocytes (PBLs) were isolated by Histopaque-1077 (Sigma, 10771) filled leucosep tubes (GBO, 227290). Total RNA was extracted from 1✕10^6^ PBLs (Qiagen, 75144) and a reverse transcription reaction was performed to produce the DNA copy (Roche, 04379012001) with oligo dT primer. The VHH sequences were amplified from the cDNA pool after a series of two PCRs with the following primers, for the first reaction: CALL001: 5’-GTCCTGGCTGCTCTTCTACAAGG-3’ and CALL002: 5’-GGTACGTGCTGTTGAACTGTTCC-3’; and for the second reaction: VHH-Back-SAPI: 5’- CTTGGCTCTTCTGTGCAGCTGCAGGAGTCTGGRGGAGG-3’ and VHH-Fw-SAPI: 5’-TGATGCTCTTCCGCTGAGGAGACGGTGACCTGGGT-3’. The Golden-Gate system and the pMECs-GG phagemid vector were used to clone Nb coding sequences between SapI sites by sequential ligation and restriction reactions [70]. Electro-competent E*. coli* TG1 cells were transformed resulting in a Nb-library of about 1.8✕10^9^ independent colonies that were resuspended in LB medium, divided into 24 tubes of 2 ml each, and stored at -80 and -196°C. Forty-eight colonies were selected and resuspended in 50 μl of H_2_O to perform a colony PCR with primers MP57: 5′- TTATGCTTCCGGCTCGTATG-3′ and GIII: 5′-CCACAGACAGCCCTCATAG-3′ in order confirm that the majority of the transformants have an insert of the proper size, as it has been suggested [51]. To rescue and amplify Nb-displaying phage particles, 2 ml of bacteria from the stock library were grown in 2xTY medium supplemented with 100 μg/ml ampicillin and 2% (w/v) glucose. When OD_600nm_=0.6 was reached bacteria were superinfected with 4✕10^10^ p.f.u. VCSM13 helper phage (Stratagene, 200251). After centrifugation to remove traces of glucose, the bacterial pellet was resuspended in 2xTY supplemented with 100 μg/ml ampicillin and 25 μg/ml kanamycin and incubated overnight at 37°C and 200 r.p.m. The next day, recombinant phages were purified by 20% (w/v) PEG 6000/2.5 M NaCl precipitation. Before the pandemic started, we had available a Nb-library obtained after immunizing a llama with 1✕10^7^ FFU/dose (fluorescent focus forming units) of the BCoV Mebus vaccine. This library was also screened with the S-2P and RBD protein from SARS-CoV-2 to detect cross-reactive Nbs.

### Isolation of SARS-CoV-2 S-2P- and RBD-specific Nanobodies

Three consecutive rounds of panning were performed as previously described [71]. Wells of a MaxiSorp microtiter plate (Nunc, 446469) coated overnight at 4°C with 0.2 μg of recombinant protein S-2P or RBD, were used to select specific binders. As negative control wells were coated with supernatants of non-transfected HEK-293T cells or non-transformed yeast, respectively. The next day, wells were washed four times with PBST (phosphate buffered saline pH 7 + 0.05 % Tween20) and blocked with 10% skimmed milk in PBST for the first round, 1% OVA in PBST for the second round and ELISA Blocker Blocking Buffer (Thermo Fisher Scientific, N502) for the third round. Approximately 1✕10^12^ phages in 100 μl of blocking solution were added and incubated for 1 h at room temperature. During the first round of panning wells were washed 10 times with PBST, while 20 washing steps were used for the second and third rounds. The remaining phages were eluted with triethylamine (TEA) 100 mM pH 10 (Sigma, T0886) for 4 min, followed by neutralization with 1 M Tris-HCl pH 8. After each round of panning, phages were amplified by infection of exponentially growing *E. coli* TG1 cells and superinfected with VCSM13 helper phage. After centrifugation to remove glucose, bacteria were grown overnight in the presence of ampicillin and kanamycin antibiotics. On the next day, phages were purified using PEG/NaCl precipitation and used for the following round of selection. Screening of specific binders was done by ELISA on 96 randomly selected individual colonies from the second and third rounds of panning, using both periplasmic extract (PE) and recombinant phages (rP). To prepare PE, individual colonies were inoculated in 1 mL of 2xTY medium containing 100 μg/ml ampicillin and 0.1% glucose in a 96-well deep well plate. After incubation for 3 h at 37°C and 200 r.p.m. Isopropyl β- D-1-thiogalactopyranoside (IPTG) at a final concentration of 1 mM was added to induce Nb expression for 4 h. Bacterial pellets were frozen and thawed twice to disrupt cells and pellets were resuspended in 100 μl of PBS. After the last step of centrifugation, the PE was collected and used for PE-ELISA (PEE). To prepare samples for rP ELISA (rPE), individual colonies were grown overnight in 0.5 ml of 2xTY medium containing 100 μg/mL ampicillin and 1% glucose. After that, 5 μl of culture from each colony was diluted in 0.5 ml of fresh medium and grown for 2 h before being infected with 5✕10^9^ VCSM13 phages/well. After 30 min of incubation without shaking, the plates were centrifuged for 15 min at 1800 g to remove glucose. Bacteria were grown overnight in 2 ml of 2xTY medium containing ampicillin and kanamycin. The next day, supernatants containing phages were pelleted by centrifugation for 15 min at 1800 g, and recombinant phages were recovered from the supernatant by PEG/NaCl precipitation.

### Periplasmic Extract ELISA (PEE) and Phage ELISA (rPE)

Nanobodies produced from PE or as rP from individual colonies were tested for binding to either SARS-CoV-2 S-2P or RBD protein. MaxiSorp microtiter plate wells (Nunc, 446469) were coated overnight at 4°C with 100 ng/well of recombinant proteins (S-2P and yRBD) or irrelevant protein as negative controls. After washing with PBST, wells were blocked with 10% skimmed milk powder in PBST and 100 μl of the PE or rP diluted ¼ was added to the wells. For PEE Nb-specific binding was detected with homemade polyclonal rabbit sera diluted 1:5000, followed by horseradish peroxidase (HRP)-linked anti-rabbit IgG diluted 1:3000 (KPL, 074-1506). In the case of rPE, HRP-labeled anti-M13 phage antibody diluted 1:3000 (GE Healthcare, RPN420) was used as a detection system. After washing, 100 μl/well of 2,2’-Azinobis [3-ethylbenzothiazoline-6-sulfonic acid]-diammonium salt (ABTS, Sigma, A1888) was added to the plates and the reaction was stopped by the addition of 5% SDS. The absorbance at 405 nm was measured using a Microplate Photometer (Multiskan™ FC, Thermo Fisher Scientific). To determine specific binding, the OD_405_ value of antigen coated wells, at least two times higher than the OD_405_ value of the control wells, were considered positive. Plasmids from positive clones were transformed in DH5α cells and further characterized by a restriction reaction with *Hinf*I before sending samples for sequencing.

### Phylogenetic analysis and germline origin

MEGA 11 was used to analyze genetic relatedness and clustering of the unique Nbs using a neighbour-joining tree with 1000 bootstrap replicates. The IMGT/V-QUEST program was used to determine the CDRs of the selected Nbs and to analyze the germline origin using their nucleotide sequences [72]. Nbs amino acid sequences were also analyzed by IMGT/DomainGapAlign [73]. Sequence logo was plotted using WebLogo3 [74].

### Expression and purification of Nanobodies

For the production and purification of the selected clones (n=43), pMECS-GG-transformed WK6 were grown at 37°C in TB medium supplemented with 100 μg/ml ampicillin and 0.1% glucose. After reaching an OD_600_ value of 0.6-0.8, Nb expression was induced with 1mM IPTG for 16 h at 28°C. Periplasmic extracts were obtained after resuspension and incubation of cell pellets with TES buffer (0.2 M Tris-HCl pH 8, 0.5 mM EDTA, 0.5 M sucrose) for 2 h, followed by incubation with TES buffer diluted 1⁄4 for 4 h. After a final centrifugation step, Nbs from PE were purified using IMAC Hi-Trap columns (GE Healthcare, 17092003) following manufacturer’s instructions and eluted using 300 nM Imidazole. Nanobodies were further purified by size exclusion gel filtration using a Superdex 75 column (GE Healthcare) pre-equilibrated with PBS using an AKTA Prime Plus Liquid Chromatography System (GE Healthcare). Afterwards, they were concentrated using Vivaspin centrifugal concentrators with a cut-off of 3 kDa (Sartorius, VS2091) and stored at -20°C. Nanobody concentration was estimated by SDS-PAGE using reference Bovine Serum Albumin Standards (Thermo Fisher Scientific, 23209) as well as by spectrophotometer using molar extinction coefficient and molecular weight (Biotek epoch 2).

### Binding affinity to the target antigen estimated by ELISA EC50

To evaluate the binding capacity of the selected Nbs to the S-2P and RBD proteins, MaxiSorp 96-well plates (Nunc, 446469) were coated overnight at 4°C with 100 ng/well of recombinant proteins in carbonate/bicarbonate buffer pH 9.6. Plates were blocked with 10% skimmed milk in PBST for 1 h at 37°C. Purified Nbs were adjusted to a concentration of 1 μM and 10-fold dilutions were added to the coated plates and incubated for 1 h at 37°C. A homemade polyclonal rabbit serum against Nbs diluted 1:2000 was added, followed by a commercial HRP-conjugated goat anti-rabbit IgG diluted 1:5000. All incubations were done at 37°C. The reaction was developed with ABTS and stopped with 5% SDS, absorbance at 405 nm was measured. The EC_50_ was estimated using a four-parameter log-logistic regression model (AAT Bioquest, Inc). Quest GraphTM IC_50_ Calculator [75].

### Pseudovirus neutralization test (pVNT)

Pseudovirus expressing Wuhan-Hu-1 SARS-CoV-2 S protein were produced by co-transfection of plasmids encoding a GFP protein (Addgene, 11619), a lentivirus backbone (VRC5602, NIH), and the S protein (VRC7475_2019-nCoV-S-WT, NIH) in HEK-293T cells as previously described [76]. To produce pseudoviruses expressing the S protein from different variants the following plasmids were used: Alpha (B.1.1.7) (InvivoGen, plv-spike-v2); Beta (B.1.351) (InvivoGEn, plv-spike-v3) and Delta (B.1.617.2) (InvivoGen, plv-spike-v8). Plasmids encoding the S protein from the Omicron variants (B.1.1.529 and BA.2) were obtained from the G2P-UK National Virology consortium.

Triplicate two-fold serial dilutions of Nbs starting at a dilution of 1 μM, or six-fold diluted heat-inactivated llama serum (56°C for 45 min) were prepared in 50 μl of OptiMem medium and combined with an equal volume of titrated pseudoviruses, incubated for 2 h at 37°C, and then added to HEK-293T cells previously transfected with plasmids coding for the ACE2 receptor and TMPRSS2 protease (VRC9260, NIH). Forty-eight hours later, cells were observed under the microscope (IX-71 OLYMPUS), and GFP-positive cells were automatically counted with ImageJ. Inhibition percentage was calculated by the following formula: 100*[1-(X-MIN)/(MAX-MIN)] where X stands for the number of GFP-positive cells at a given concentration of Nb and MIN and MAX refers to the number of GFP-positive cells in uninfected cells or in cells transduced with only pseudovirus respectively. IC_50_ titers were determined based on sigmoidal nonlinear regression using GraphPad Prism software Inc (La Jolla, CA, USA) [77].

### Wild type virus neutralization assays (VNA)

Neutralization assays were done at the Virology Institute, INTA using the isolate hCoV- 19/Argentina/PAIS-C0102/2020 (D614G). Vero cells, 1.5✕10^4^ cells/well were seeded in 96-well plates and incubated for 24 h at 37°C in 5% CO_2_ atmosphere. Two-fold diluted Nbs (starting concentration 10 μM), were incubated with 100 TCID_50_ of the virus at 37°C for 1 h in DMEM supplemented with 2% fetal bovine serum, penicillin/streptomycin and 10 μg/ml amphotericin B. Cells were infected with virus/Nb mixture and incubated for 1 h at 37°C and 5% CO_2_. Inoculum was washed and cells were incubated with D-MEM with 2% fetal bovine serum for 72 h at 37°C until a cytopathic effect (CPE) was observed. Cells were then fixed with 70% acetone. Virus replication was confirmed by immunofluorescent staining using a homemade FITC-labeled polyclonal IgG llama serum produced against SARS-CoV-2 S-2P protein. To produce this reagent, llama IgG was purified from serum by ammonium sulfate precipitation, dialyzed against carbonate-bicarbonate buffer (pH 9.0) and labeled with fluorescein isothiocyanate (FITC, Sigma F3651). The fluorescein-labeled antibody was separated from the unconjugated antibodies by Sephadex G-25M size exclusion chromatography. The F/P molar ratio of the purified protein was determined by measuring the absorbance at 280 nm and 495 nm. Cells were stained by adding 50 μl of the FICT-labeled polyclonal antibodies per well at a 1:100 dilution in PBS with Evans blue and read under a fluorescence microscope. The neutralizing titer was calculated as the inverse of the highest dilution that evidences positive fluorescence, comparable to non-infected Vero cells.

A plaque reduction neutralization test (PRNT) was performed using a United States isolated (USA- WA1/2020, NR-52281). Two-fold serially diluted Nbs in DMEM medium supplemented with 2% FBS (starting concentration 1 μM), were incubated with a viral suspension containing 100 plaque-forming units of SARS-CoV-2 virus at 37°C for 72 h. Cells were then fixed with 4% paraformaldehyde for 20 min at 4°C and stained with crystal violet solution in methanol. The CPE was assessed visually considering minor damage to the monolayer (1–2 plaques) as well as a manifestation of CPE. Neutralization titer was defined as the highest serum dilution without any CPE in two of three replicable wells. The number of infected cells was determined per well by counting localized areas of clearance in the cell monolayer left undeveloped by the crystal violet. PRNT_50_ was calculated using the NIAID Calculator [78]. The maximal inhibitory concentration was established at 90% (IC_90_) reduction of the CPE detected at light microscope corresponding also to the same reduction of fluorescent focus forming units.

### Competition of Nanobodies with ACE2

To determine whether the Nbs prevent the SARS-CoV-2-RBD/ACE2 interaction, we set up a surrogate virus neutralization test based on the ELISA technique. For this, RBD was adsorbed overnight at 4°C in 96-well plates at a concentration of 0.2 μg/well in carbonate/bicarbonate buffer pH 9.6. Plates were washed 3 times with PBST and blocked for 1 h at RT with 200 μl 3% skimmed milk. Afterward, two-fold serial dilutions of specific and irrelevant Nbs (starting concentration 1 μM) were added to the wells and incubated for 1 h at room temperature. Recombinant ACE2-HRP produced in HEK-293T cells was incubated for 2 h at RT. Wells were washed 3 times with PBST, and 50 μl of 3,3’, 5,5’ tetramethylbenzidine (TMB, BD, 555214) was added for 15 min. The reaction was stopped with 50 μl H_2_SO_4_ and the plates were read in a spectrophotometer at 450 nm (TECAN, Infinite 200). The binding kinetics was analyzed by a non-linear regression model using GraphPad Prism software Inc (La Jolla, CA, USA).

### Interference of Nanobody binding to Spike by biliverdin

Nanobodies that were selected with the S-2P protein and do not bind to RBD, were tested in an ELISA-based biliverdin competition assay. For this, biliverdin IX α (biliverdin) was obtained by oxidative ring opening of hemin IX (Sigma Aldrich, 51280) following a reported protocol [79,80]. Esterification of the material was performed by dissolution in BF_3_/CH_3_OH. The green solution was kept for 12 h in the dark, and then diluted with water and extracted with CH_2_Cl_2_, the organic phase was washed with water, dried over Na_2_SO_4_, filtered and evaporated to dryness. The crude mixture of dimethyl esters was purified by TLC chromatography [81]. After silica removal, the separated dimethyl esters were recrystallized from CH_2_Cl_2_:CH_3_OH. The separated blue-green pigments (biliverdins IX α, β, γ and δ) were characterized by NMR. Pure biliverdin IX α was saponified by dissolution in CH_3_OH under an argon atmosphere, followed by the addition of an equal volume of 1M NaOH containing 1 mM EDTA. The solution was kept in the dark for 2 h, and then it was poured over water and extracted into CH_2_Cl_2_ with the addition of CH₃COOH. The blue-green organic phase was washed with water, dried over Na_2_SO_4_, filtered and evaporated to dryness. Stock solution was prepared in DMSO and diluted to 1% for working solutions. Concentration was determined by absorbance at 388nm (ε= 39900 M^-1^cm^-1^).

Ninety-six-well plates were coated overnight with 0.05 µg of purified S-2P protein and next day blocked with 1% skimmed milk in 0.5% PBST. Subsequently, 25 µl of 10 µM biliverdin was added to half of the wells and incubated for 5 min. Then 25 µl of ten-fold diluted Nbs were added (starting concentration 1 µM) and incubated for 1 h at RT. Specific binding was determined using an HRP-conjugated anti-HA antibody diluted 1:5000 (Abcam, Ab1190), and the reaction was developed with TMB substrate and read at 450 nm. In a second experiment, a fixed concentration of Nbs (0.1 µM) was mixed with a five-fold serial dilution of biliverdin (starting concentration 25 µM) and the reaction was developed as previously mentioned.

### Western Blot analysis

The reactivity of each Nb was analyzed by Western blot (WB). For this assay, 0.5 μg of recombinant S-2P or RBD proteins were separated in 8-10% SDS-PAGE under three different conditions: a) reducing and denaturing conditions, b) non-reducing but denaturing conditions, and c) non-reducing and non-denaturing conditions. The proteins were blotted onto a nitrocellulose membrane using a semi-dry transfer system (Bio-Rad). The membranes were blocked with 5% skimmed milk in PBS for 1 h at RT. Each Nb (1 μg/ml) was incubated with the transferred proteins for 2 h and the binding was detected after incubation for 1 h at RT with HRP-conjugated anti-HA antibody diluted 1:5000 (Abcam, Ab1190). The binding of the conjugate was detected with ECL Western Blotting Substrate (Thermo Fisher Scientific, 32106).

### Efficacy in a mouse model

To assess the protective efficacy of the Nbs against SARS-CoV-2 infection, 4-week-old k18-hACE2 mice (Jackson Labs) were separated into 7 groups (n=9) of mice with approximately equal numbers of males and females in each group. Approximately 4 h before challenge, mice were administered 10-20 µg of anti-rotavirus control Nb (2KD1), anti-SARS-CoV-2 Nb-33, Nb-39, Nb-43, Nb-45, Nb-104, or Nb-110. Nanobody 145 was not tested in the mouse model due to the low availability of animals and the high similarity in the CDR3 sequence of this Nb and Nb-39. Mice were then challenged intranasally with 1✕10^5^ PFU of the WA1/2020 strain of SARS-CoV-2 in each nostril. Mice were then monitored daily for weight loss and survival, with checks increasing to at least 3 times daily when disease symptoms presented. Four days post-challenge, 3 mice in each group (1 male, 2 females, excluded from weight and survival data) were euthanized, and tissues (i.e. brain, lungs and nasal turbinates) were collected to assess the impact of the Nb treatment on viral titers by RT-qPCR as previously described [82]. Primer sequences were taken from 2019-Novel Coronavirus (2019-nCoV) Real-time rRT-PCR Panel from the Centers for Disease Control and Prevention (CDC). Quantitative synthetic SARS-CoV- 2 RNA from ORF1ab, E and N was used for the generation of a standard curve to determine viral load (ATCC, VR-3276SD). The survival data were analyzed by the Mantel-Cox log-rank test. Virus titers were log10 transformed and analyzed under a general linear mixed model analysis of variance, where treatment and tissue were considered fixed factors with interaction. The heterogeneity of variance among groups was modeled using a varIdent variance-covariance matrix. Post ANOVA multiple comparisons of the mean virus load in each treatment group (overall mean) and in each tissue inside each treatment group were analyzed by LSD Fisher test and p-values corrected by the Bonferroni method. The analyses were conducted in Infostat with a link to R [83]. All animal experiments and operations were performed in the biosafety level 3 (BSL-3) facility, and the protocols were approved by the Institutional Animal Care and Use Committee at Virginia Tech, IACUC number 21-065 SARS-CoV-2, Pathogenesis and Countermeasure Testing.

### Epitope mapping of Nanobodies

To determine binding epitopes for Nbs in study, a 181-peptide array covering the entire amino acid sequence of the SARS-CoV-2 S glycoprotein was used (BEI resources, NR-52402). Peptides were resuspended in 500 μl of DMSO:PBS to obtain a 2 mg/ml stock. Maxisorp 96-well plates were coated with 100 ng of each peptide in carbonate/bicarbonate buffer pH 9.6 and incubated overnight at 4°C. After 3 washing steps with PBST and a blocking step using Starting Block PBS (Thermo Fisher Scientific, PI37538) for 1 h at room temperature, Nbs (100 ng/well) were added for 2 h at room temperature. The positive reaction was developed by polyclonal homemade rabbit serum against Nbs diluted 1:5000 and HRP-conjugated goat anti-rabbit IgG diluted 1:5000 or anti-6xHis-HRP. After washing the reaction was developed with TMB and read at 450 nm.

### In silico analysis

#### Selection of the Spike protein and Nanobodies structures

The trimeric form of the Spike protein of coronavirus (pdbID 6vyb) was selected for performing protein-protein docking experiments with the Nb-43, Nb-45, and Nb-53. These Nbs were selected based on their high affinity for the S protein and their ability to enhance the neutralization potency in a Nb cocktail.

#### Nanobody modeling and Molecular dynamic refinement

Nanobody 43, Nb-45 and Nb-53 were modeled using deep learning-based end-to-end modeling, Nanonet [84]. The top-rated structures were selected and subsequently refined for 100 ns using molecular dynamics simulation with AMBER22. To maintain homogeneity, identical parameters were used for all cases. First, each system was optimized using a conjugate gradient algorithm for 5000 steps. This was followed by two thermalizations: a 10 ps constant-volume MD at 10 K with a restraint of 50 kcal/mol, and a second thermalization in which the temperature of the system was gradually increased from 10 to 300 K over 100 ps (integration step = 0.0005 ps/step). The system was then equilibrated for 250 ps at constant temperature and pressure (integration step = 0.001 ps/step) to reach the desired system density. A second equilibration MD of 500 ps was performed with an increased integration step of 2 fs and a decreased force constant for restrained alpha-carbons of 2 kcal/mol/Å. Finally, a 10 ns long MD simulation was performed without constraints using the “Hydrogen Mass Repartition’’ technique which allowed for an integration step of 4 fs [85]. These conditions were used for all subsequent production of 1 us long MD runs with 10 steps per ns and 5,000 steps per simulation. The Amber package of programs was used for all simulations, using the ff19SB force field for all AA residues and the

OPCBOX model for water molecules as solvent [86]. Monte-Carlo barostat and Langevin thermostat were used to maintain constant pressure and temperature, respectively, with default coupling parameters. A 10 Å cut-off for non-bonded interactions was used, and periodic boundary conditions were applied using the Particle Mesh Ewald summation method for long-range electrostatic interactions. VMD software was used for visualizations and image rendering.

#### High Ambiguity Driven protein-protein Docking

Protein-protein docking was performed using the High Ambiguity Driven protein-protein Docking (HADDOCK) with a semi-flexible and unrestricted docking protocol [87,88]. The prefused Spike RBD up (6VYB) was used as the docking template, and multiple docking runs were performed with Nb-43, Nb-45, and Nb-53 against the S protein to identify the best result based on the lowest RMSD clustering score. Additionally, docking of the ligand biliverdin to a specific region of the S protein (AA 121-207) was carried out to evaluate its biological relevance. Electrostatic calculations were performed using a distance-dependent dielectric, which was chosen based on its suitability for studying protein-ligand interactions. No water refinement was applied.

#### Protein-protein docking prediction

Protein-protein docking experiments were performed using the HADDOCK 2.4 server. Three different results were generated for each combination of Nb and chain of the S trimer. For each result, the best option was selected based on the best score and the best pose. In addition, it was taken into account that Nbs bind to the S protein trimer mostly using the CDRs.

#### Protein-ligand docking with biliverdin

A protein-ligand docking experiment was also performed with biliverdin using the HADDOCK 2.4 server. The specific region of the S protein where biliverdin binds was identified based on the AA reported in the literature [27]. The protein-ligand docking was performed on this identified region to evaluate the biological relevance of the docking results.

#### Interaction analysis

To analyze the molecular interactions among the docking results, a custom Python script was developed. The script calculates the distance between the AA of the Nb and the S protein by computing the center of mass of each residue within a 7 Å distance. This generates a list of potential interacting residues. To analyze these interactions, the list of residues was imported into VMD, and the distance tool was used to determine the distance between specific AA. The knowledge of residue interactions was then used to infer which AA were likely interacting between the Nb and the S protein. The types of AA interactions were also identified and analyzed.

## Acknowledgments

We thank Stephan Kissler for the pLB plasmid (Addgene, 11619). We express gratitude to Dr. Karin Bok (VRC, NIH) for the SARS-CoV-2 S-2P, RBD, ACE2_TM, TMPRSS2 plasmids and to Dr. Florian Krammer (ISMMS) for the SARS-CoV-2 S-2P and RBD plasmids. Dr. Yves Durocher, NRCC, Canada for the S protein expressed in CHO cells. The following reagents were obtained through BEI Resources, NIAID, NIH: Peptide Array, SARS-Related Coronavirus 2 Spike (S) Glycoprotein, NR-52402; SARS-Related Coronavirus 2, Isolate hCoV-19/USA-WA1/2020, NR-52281. We acknowledge the G2P-UK National Virology consortium funded by MRC/UKRI (grant ref: MR/W005611/1.) and the Barclay Lab at Imperial College for providing the Omicron B.1.1.529 and Omicron BA.2 spike-encoding plasmids. Special thanks to “Cabaña Las Lilas S.A.” for providing the llamas for free during the hardest time of the pandemic and Mr. Aimar for his help in their transportation. Thanks to Dr. Matias Aduriz for the BCoV library. The Argentinian AntiCovid Consortium is formed by Amante Analía, Blaustein Matías, Bredeston Luis, Corapi Enrique, Craig Patricio, D’Alessio Cecilia, Elias Fernanda, Fernández Natalia Brenda, Gándola Yamila, Gasulla Javier, Gentili Hernan, Gorojovsky Natalia, Gudesblat Gustavo, Herrera María Georgina, Ibanez Itatí, Idrovo Tommy, Iglesias Matias, Kamenetzky Laura, Nadra Alejandro, Noseda Diego, Pavan Carlos, Pavan Florencia, Pignataro María Florencia, Roman Ernesto, Ruberto Lucas, Rubinstein Natalia, Santos Javier, Velazquez Duarte Francisco, Wetzler Diana and Zelada Alicia.

## Supporting Information

**S1 Fig.**
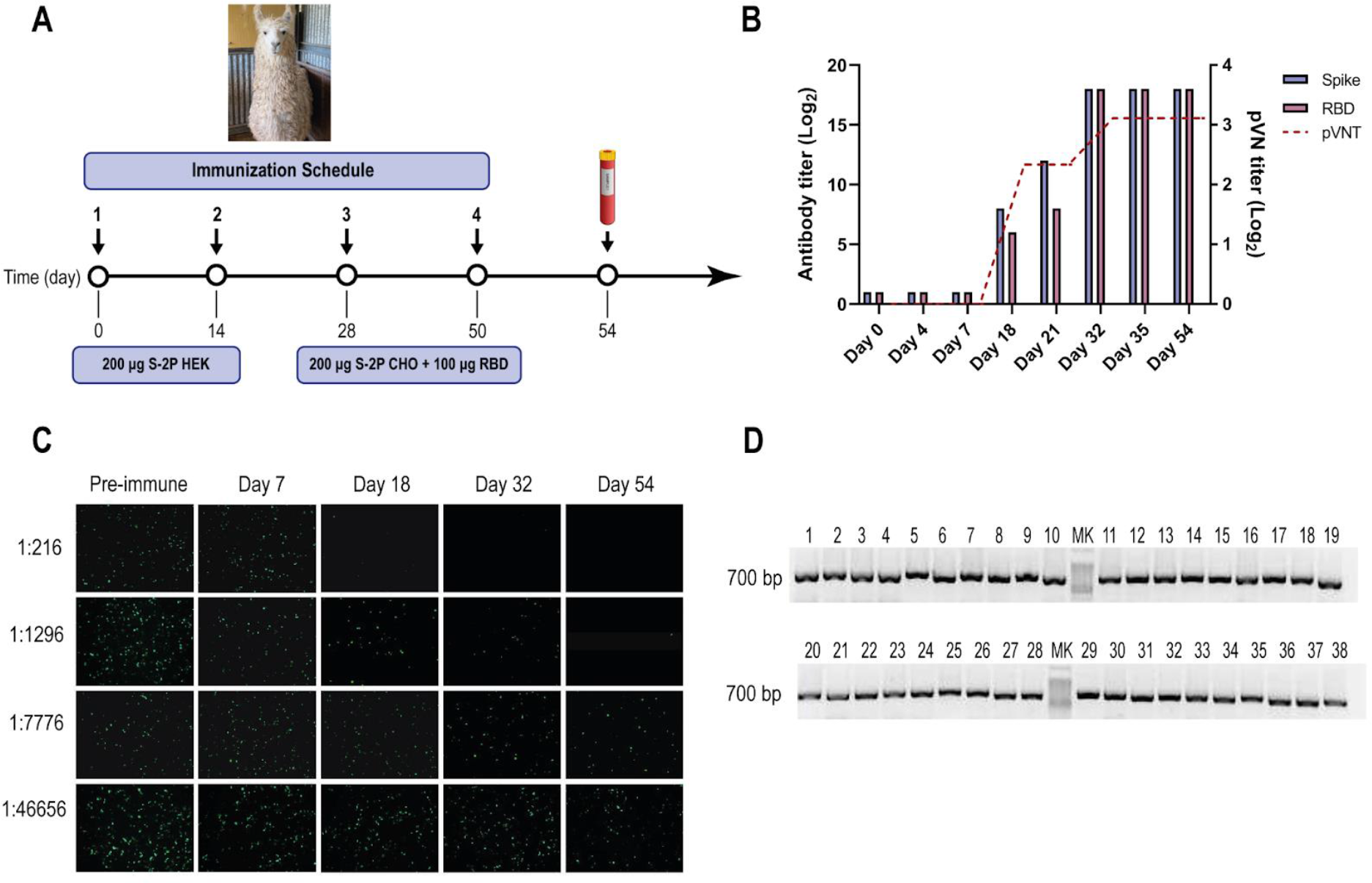
SARS-CoV-2 llama immunization, immune response and Nb-library construction. (**A**) Immunization schedule: a llama was injected intramuscularly on days 0 and 14 with 200 μg of SARS-CoV-2 S-2P protein produced in HEK-293T and on days 28 and 56 with 200 μg of SARS-CoV-2 S-2P produced in CHO cells and 100 μg of RBD protein emulsified in Freund’s adjuvant. Four days after the last boost, 200 ml of blood was collected, and peripheral lymphocytes were isolated to produce an immune library. (**B**) Total IgG titer determined by ELISA and neutralizing Ab titer determined by pVNT induced 4 and 7 days after each immunization. Four days after the third immunization (PID 32) a maximal antibody response was reached. (**C**) Picture illustrating neutralizing activity in llama serum determined by pVNT. The neutralization capacity increased after each immunization and correlated with a decrease in the number of fluorescent cells. A higher neutralizing titer was detected for a dilution of 1:1296 at PID 54. (**D**) Analysis of PCR products by agarose gel electrophoresis to confirm the number of transformants that had an insert of the proper size, each of the 48 clones that were randomly selected contained a genuine Nb fragment (∼700 bp).

**S2 Fig.**
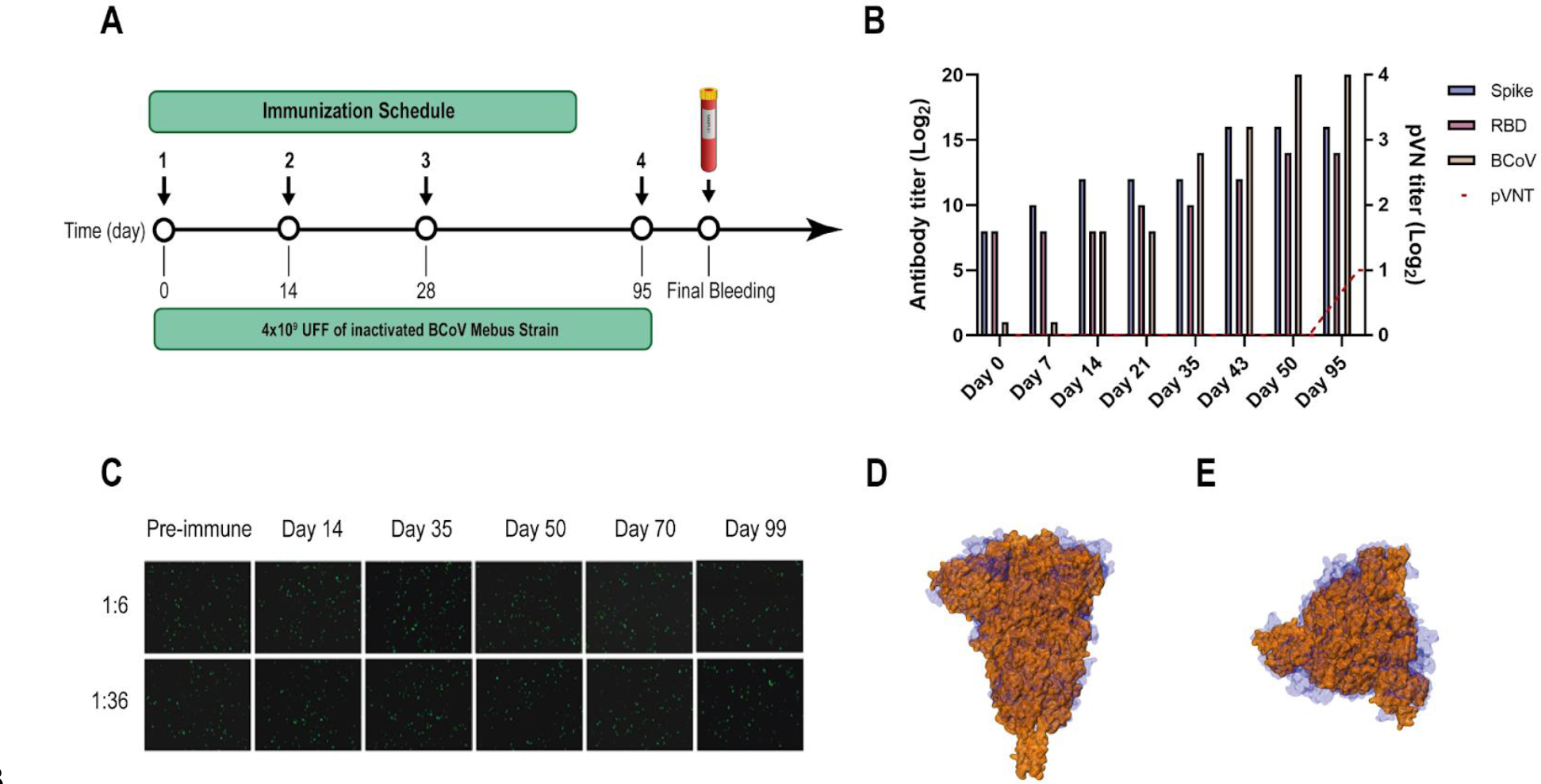
**BCoV Mebus llama immunization and immune response**. (**A**) Immunization schedule followed to produce the Nb immune library: a llama was injected intramuscularly on days 0, 14, 28 and 95 with 4.00
✕109 UFF of the inactivated BCoV Mebus strain in Freund’s adjuvant. Peripheral lymphocytes were isolated from 200 ml of blood collected four days after the final boost to generate the immune library. (**B**) Total IgG titer determined by ELISA for BCoV and SARS-CoV-2 RBD and S-2P proteins. (**C**) Picture showing non-neutralizing activity of sera from a llama immunized with BCoV Mebus against SARS-CoV-2 determined by pVNT. Superimposition of the S protein structures from SARS-CoV-2 (orange) and BCoV Mebus (blue) using VMD software. Frontal (**D**) and upper (**E**) view.

**S3 Fig.**
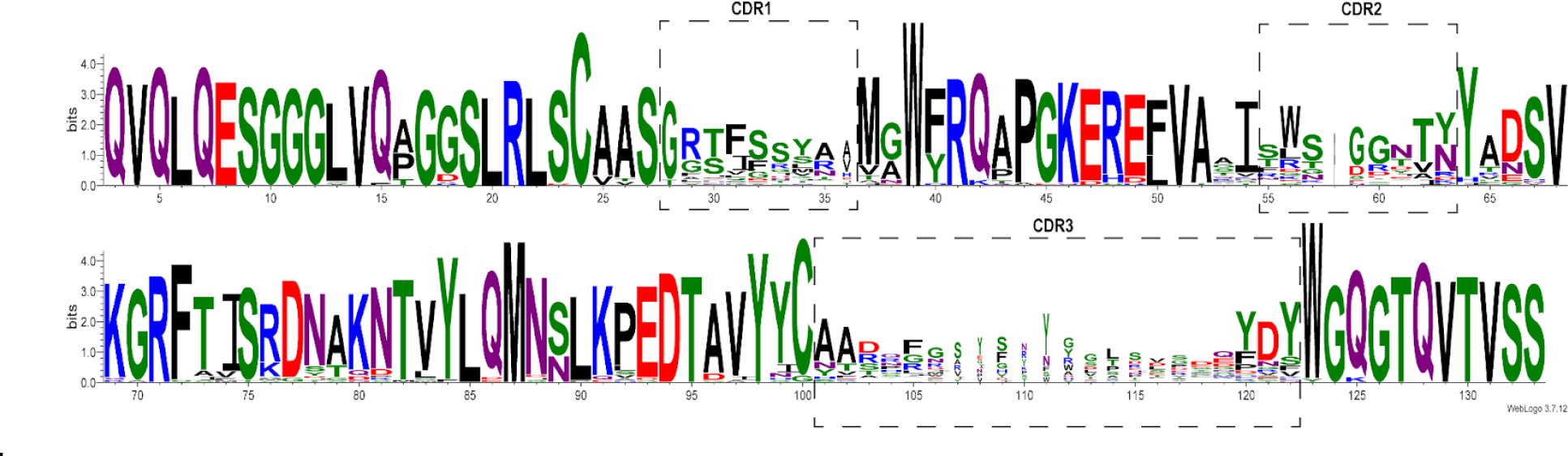
**Sequence Logo Plot of a diverse repertoire of SARS-CoV-2 Nbs**. Logo representation of amino acid multiple sequence alignments of the 43 unique Nbs selected after biopanning with RBD or S-2P proteins. The height of symbols indicates the relative frequency of each amino acid at that position.

**S4 Fig.**
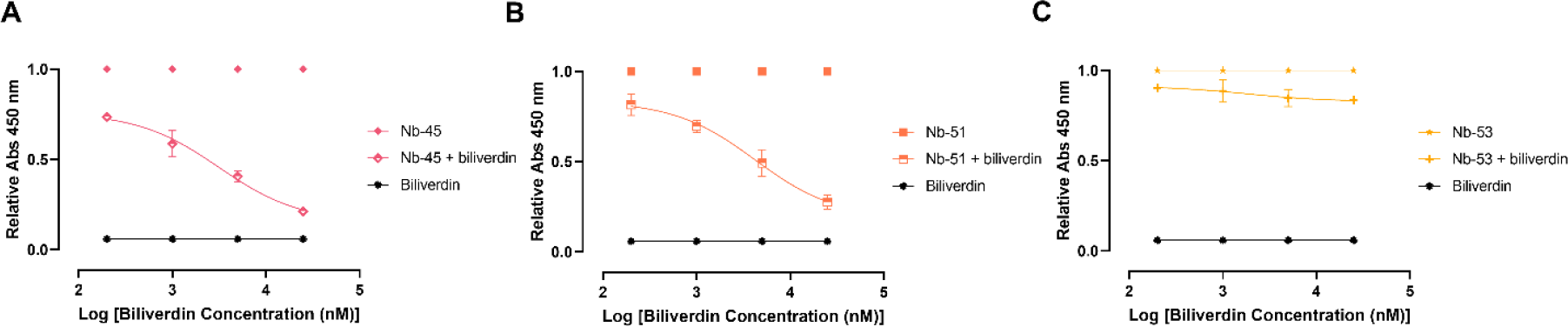
**Biliverdin decreases the binding of non-RBD binders to S-2P protein**. Relative dose-response curves for Nb-45 (**A**), Nb-51 (**B**) and Nb-53 (**C**) in the absence or presence of different concentrations of biliverdin.

**S5 Fig.**
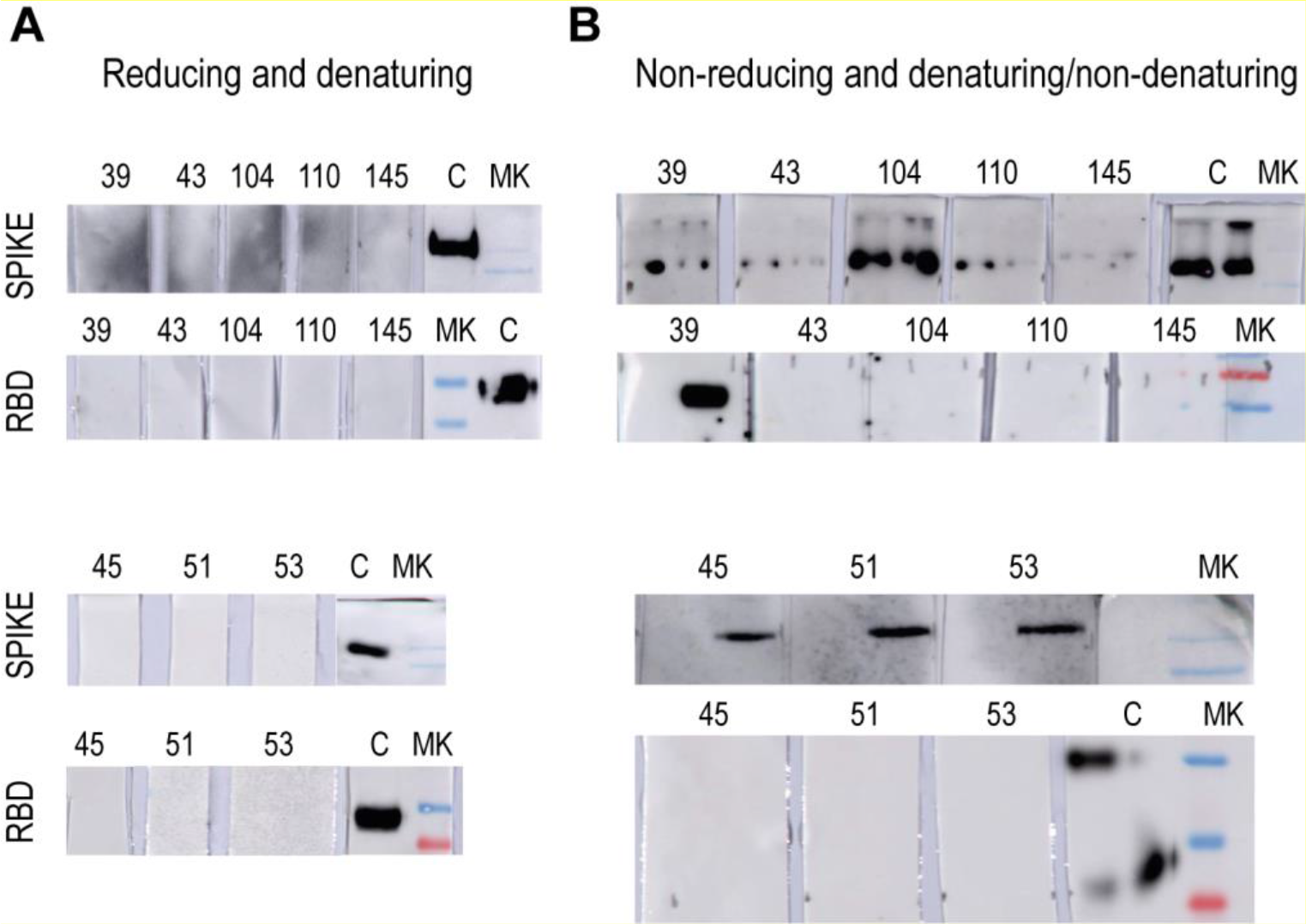
Determination of Nanobodies binding to conformational or lineal epitopes by Western blot analysis. Different conditions of sample buffer and gel composition were tested to determine the Nbs reactivity. (**A**) Reactivity of Nbs to S-2P and RBD proteins under reducing and denaturing conditions. (**B**) Nanobodies recognition of S-2P and RBD proteins under non-reducing and denaturing (left well) or non-denaturing conditions (right well) for each Nbs. As can be observed, Nb-43, Nb-104, Nb-110 and Nb-145 can detect the S-2P protein only under non-denaturing conditions while Nb-39 recognizes not only S-2P but also RBD. Nanobody 45, Nb-51 and Nb-53 detect only the S-2P protein under non-denaturing conditions and do not bind to RBD.

**S1 Table.**
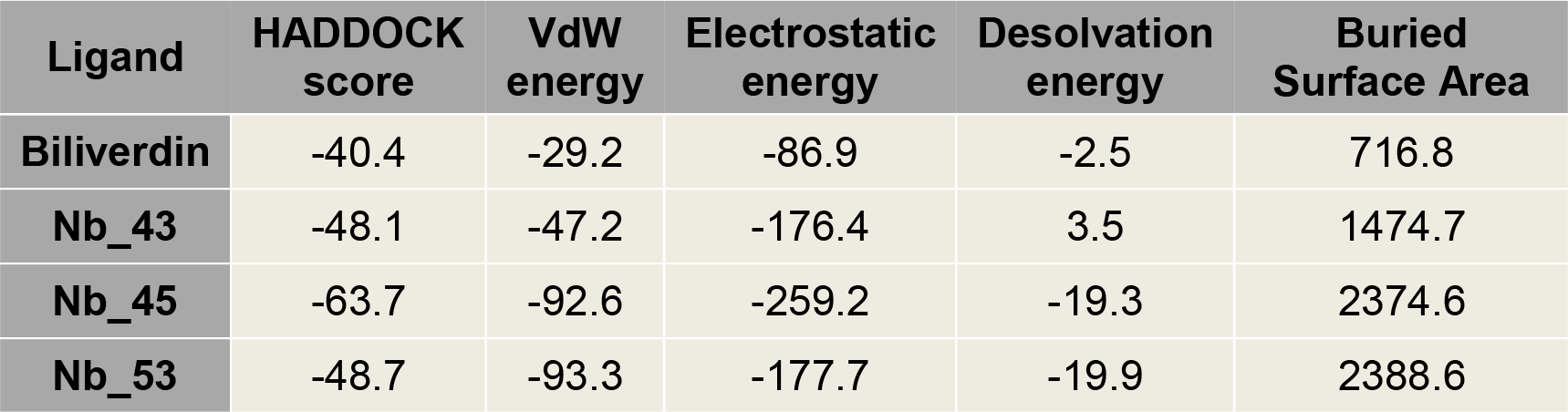
Summary of HADDOCK results for the best Nanobody-Protein complexes.

| Ligand | HADDOCK score | VdW energy | Electrostatic energy | Desolvation energy | Buried Surface Area |
| --- | --- | --- | --- | --- | --- |
| Biliverdin | -40.4 | -29.2 | -86.9 | -2.5 | 716.8 |
| Nb_43 | -48.1 | -47.2 | -176.4 | 3.5 | 1474.7 |
| Nb_45 | -63.7 | -92.6 | -259.2 | -19.3 | 2374.6 |
| Nb_53 | -48.7 | -93.3 | -177.7 | -19.9 | 2388.6 |

